# Hybrid crosses reveal a cell-type-specific landscape of mouse regulatory variation

**DOI:** 10.64898/2026.04.02.716195

**Authors:** Ryan Weber, Maria Carilli, Elisabeth Rebboah, Ghassan Filimban, Heidi Yahan Liang, Diane Trout, Margaret Duffield, Parvin Mahdipoor, Erisa Taghizadeh, Negar Fattahi, Negar Mojgani, Romina Mojaverzargar, Nikhila Swarna, Shimako Kawauchi, Brian A. Williams, Grant R. MacGregor, Barbara J. Wold, Lior Pachter, Ingileif B. Hallgrimsdottir, Ali Mortazavi

## Abstract

Understanding the genetic architecture of gene expression is fundamental to evolutionary biology and medicine. As part of the IGVF Consortium, we present a single-nucleus RNA-seq resource of 5.3 million nuclei across eight tissue groups, featuring seven F1 hybrids from C57BL/6J dams crossed with the other Collaborative Cross founder strains for comparison against parental strains. We identify 25,864 genes (91% of those detected) exhibiting non-conserved regulatory behavior in at least one of 92 cell types in one or more crosses. Our results show that while cis-acting variation primarily drives divergence, trans-acting effects are substantially more cell-type specific and sensitive to tissue environment. Notably, bulk tissue analyses frequently mask these signals, particularly in smaller populations such as astrocytes. Furthermore, increasing genetic divergence primarily expands the landscape of cis-acting variation, while trans-acting effects remain stable across genetic distances within species. This atlas establishes a foundational framework for decoding the complex interplay between genetic variation and cell-type-specific regulation across the mammalian body.

## Introduction

Heritable non-coding variants account for a substantial portion of the genetic contribution to phenotypic diversity, arising from effects of common and rare variants in non-coding regulatory elements^1^. However, how changes in genetic architecture alter gene expression, and ultimately influence traits, remains poorly understood^2^. Advances in single-cell genomics enable the characterization of how genetic variation shapes gene expression diversity across distinct cell populations in a cell-type-specific manner^3^. A central goal of the Impact of Genomic Variation on Function (IGVF) consortium is to link genomic variation to function and phenotype by understanding how non-coding variation influences gene expression and downstream phenotypes in both humans and mice^4^.

Gene expression variability attributed to genetic variation can be explained by cis-acting and/or trans-acting genetic variants^5^. Specifically, cis-acting variants can impact a gene’s expression by altering sequences at or near motifs within nearby cis-regulatory elements that regulate the target gene^6^. Cis-acting variants that modulate the expression of diffusible trans-regulatory elements (such as transcrip-tion factors) can, in turn, influence the expression of many downstream target genes, thereby giving rise to potentially pleiotropic trans-acting effects^7,8^. Accordingly, variation in the expression of a single gene can reflect the combined influence of both cis-acting and trans-acting variants^9^. When these effects act in the same direction, they contribute additively to gene expression (cis+trans-acting). In contrast, when cis-and-trans-acting variants act in opposing directions (cis *×* trans-acting), their effects partially or fully counteract one another; such interactions may also be referred to as compensatory regulation^10^.

One approach that has been developed to measure the contribution of cis- and trans-acting variation to gene expression variability is by comparing the ratio of gene expression between homozygous parents to the ratio of allele-specific gene expression in F1 hybrids^11,12^. Under this framework, genes whose allele-specific expression differences in F1 hybrids match the expression differences observed between the two homozygous parents are inferred to be driven primarily by cis-acting variation. In contrast, genes that show expression differences between parents but no allele-specific imbalance in F1 hybrids are consistent with regulation by transacting variation. Analyses applying this approach have revealed substantial variation in the relative contributions of cis-, trans-, cis+trans-, and cis*×* trans-acting regulation, and have been applied across a wide range of organisms, including yeast,^13^ flies^14^, sticklebacks^15^, mice^16^, and plants^17^.

Previous applications of this framework in mice have provided critical insights into the regulatory landscape across diverse tissues, including liver^16^, retina^18^, and testes^19^, and have characterized how regulatory effects shift in response to environmental changes^20–22^. Together, these studies quantified cis-acting and trans-acting regulatory effects across defined mouse genetic backgrounds and tissues, establishing patterns of regulatory variation at the tissue level. Because these approaches relied on bulk tissue assays, they capture gene expression as an aggregate signal across heterogeneous cell populations. As a result, the extent to which cis- and trans-acting regulatory variation operates at cell-type-specific resolution and how these patterns generalize across diverse genetic backgrounds remains largely unexplored in the mouse.

A powerful resource for enabling genetic diversity studies in a controlled system is the recombinant inbred Collaborative Cross (CC) mouse panel^23^. The eight CC founder mice include five laboratory-derived strains, including the most commonly used laboratory strain C57BL/6J (B6J), A/J (AJ), NOD/ShiLtJ (NOD), 129S1/SvImJ (129S1), and NZO/HlLtJ (NZO), as well as three wild-derived strains: WSB/EiJ (WSB), PWK/PhJ (PWK), and CAST/EiJ (CAST).Together, these eight CC founder mice capture over 90% of the genetic diversity present in *Mus musculus*^24–26^. We have previously investigated gene-expression variation across the Collaborative Cross founder strains by generating a large single-nucleus RNA-sequencing (snRNA-seq) dataset spanning eight core tissues and all eight founders, identifying widespread, cell-type-specific transcriptional differences dependent on genetic background^25^.

Expanding on this map of genetic diversity, we present a snRNA-seq resource of seven F1 hybrid mice generated by crossing B6J dams with the other seven CC founder mice. By integrating these data with our existing founder dataset, we enable a high-resolution dissection of the cis- and transacting regulatory variation underlying expression differences across strains. This consolidated atlas spans 15 genotypes, 8 core tissues, and 92 distinct cell types and states. Our analysis reveals widespread, cell-type-specific regulatory architecture that is highly dependent on both tissue context and genetic background.

## Results

### A multi-tissue single-cell resource enables cis- and trans-regulatory inference

We generated a snRNA-seq dataset from F1 mice using combinatorial split-pool barcoding, designed for integrative analysis with our existing CC founder dataset (Rebboah et al., 2025) to quantify cis- and trans-acting regulatory variation (Fig. 1A,B). Whereas the founder dataset captures gene expression variation across inbred parental strains, the F1 dataset enables allele-specific expression measurements required to disentangle cis- and transacting effects. The F1 dataset comprises seven genotypes generated by crossing B6J dams with sires from each of the remaining seven CC founder strains, along with eight additional B6J samples that allow us to compare to the original experiment (Fig. S1A). In contrast to the founder dataset, in which each tissue was distributed across two split-pool barcoding plates, the F1 dataset was generated with one tissue per plate (Fig. S1B). A total of eight combinatorial barcoding plates were used to generate the dataset, sequenced with 177 billion short reads across 8 million nominal nuclei.

**Figure 1.**
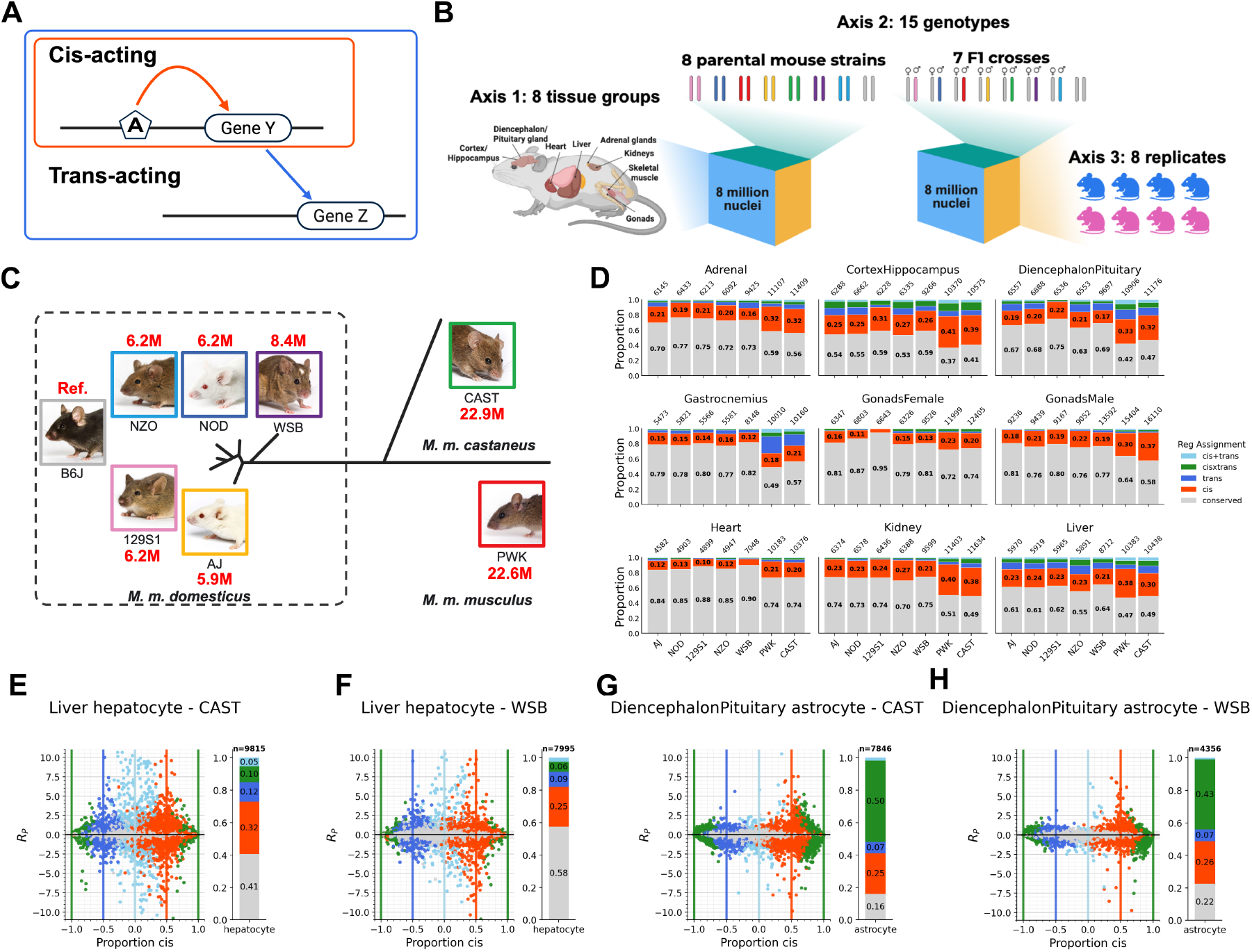
A multi-tissue, multi-strain single-nucleus RNA-seq dataset for inferring regulatory variation. (A) Schematic representation of cis-acting and trans-acting genetic variants. (B) Overview of the dataset across eight tissue groups, 15 genotypes, and eight replicates. (C) Genetic variation present among the eight CC founder strains. (D) Bulk whole-tissue regulatory assignments in seven parental-F1 trios. (E–H) Log_2_ fold change in expression between parents (*R*_*P*_ ) by proportion cis for each gene in a cell type. Stacked bar plots depict the breakdown of regulatory assignments for each cell type: (E) liver hepatocytes in B6J-CAST trios, (F) liver hepatocytes in B6J-WSB trios, (G) diencephalon/pituitary astrocytes in B6J-CAST trios, and (H) diencephalon/pituitary astrocytes in B6J-WSB trios.

Following ambient RNA removal with CellBender^27^ and quality control filtering, we recovered 5,346,886 nuclei. Across all eight genotypes and replicates, we recovered 780,686 adrenal gland, 613,914 isocortex and hippocampus (hereafter referred to as the cortex/hippocampus), 898,485 diencephalon/pituitary gland, 812,051 gastrocnemius, 549,058 female gonad, 309,543 male gonad, 95,425 heart, 707,040 kidney, and 580,684 liver nuclei. When integrated with our previously generated CC founder dataset, the combined dataset spans 637,460 to 1,863,454 nuclei per tissue (Fig. S1C).

Allele-specific expression in F1 mice can be measured for genes that contain single nucleotide polymorphisms (SNPs) on the non-B6J allele. The eight CC founder parental mouse strains are comprised of five laboratory strains (B6J, AJ, NOD, 129S1, and NZO) and three wild-derived strains (WSB, PWK, and CAST), and contain 5.9–22.9 million SNPs compared to the B6J reference mouse strain (Fig. 1C)^28^. Across the seven non-B6J founder strains, 60–90% of genes contain at least one SNP relative to B6J, with interstrain variation (Fig. S1D). More divergent strains, particularly PWK and CAST, contain SNPs in nearly all proteincoding genes, providing broad power for allele-specific expression quantification.

We recorded both total body weight (Fig. S2A) and the weight of each of the nine tissues (Fig. S2B-J) for each replicate of the F1 mouse genotypes and the repeat B6J mice and compared them to the weights of the parental strain tissues in the founder strains^25^. B6J-NZO and B6J-WSB F1 mice exhibited intermediate body weights compared with both parents. Both B6J-PWK F1 and PWK mice had smaller body weights compared with B6J, and CAST mice had smaller body weights than either B6J or their F1. Notably, we observed that female gonads of F1 mice were significantly heavier than either parent, with the exception of B6J-AJ F1 mice, which showed no difference in female gonad weight (Fig. S2F). Broadly, these results reveal that the inheritance of tissue mass is not uniform. While the liver demonstrates additive phenotypes, other tissues, notably the male and female gonads and gastrocnemius, exhibit non-additive effects, with weights in the F1 mice exceeding those of either parent. Overall, these results demonstrate a range of both additive and non-additive weight phenotypes present between parental and F1 mouse genotypes.

To quantify allele-specific expression in each of the seven F1 crosses, we generated a reference genome containing both the B6J and non-B6J alleles for each respective cross. Briefly, we generated a strain-specific version of the mm39 reference genome by swapping nucleotides overlapping SNPs for the non-reference allele using g2gtools (see Methods). The strain-specific reference genome was then concatenated with the standard mm39 reference genome to generate a reference containing both alleles for a given F1 cross. For each parental-F1 trio, we mapped both parents and the F1 to the concatenated reference using kb-python^29^ to quantify allele-specific counts. We performed quality control and cell-type annotation using single-genome mappings to mm39 in order to maximize read mapping and ensure a standardized pipeline across all genetic backgrounds. Using these single-genome mappings, we annotated 92 cell types and states, 14 fewer than previously reported in Rebboah et al., 2025, due to the consolidation of closely related cell states. For convenient exploration of results, a containerized, interactive data viewer can be downloaded from https://github.com/mortazavilab/mousaic.

### Cell-type specificity of regulatory composition

Using strain-specific mappings for the parental strains and allele-specific mappings for the F1 hybrids, we estimated cis-acting and trans-acting contributions to gene expression variability using the XgeneR package^30^. This framework utilizes generalized linear models (GLMs) to quantify regulatory differences between parental strains using parental and hybrid allele-specific expression data. XgeneR fits a negative binomial GLM to both parental expression and hybrid allele-specific expression, with weights modeling cis- and transacting regulatory differences. It then tests whether these weights are significantly different from zero using likelihood ratio tests, and genes are classified as being regulated differently in parental strains primarily as cis-acting, trans-acting, or a combination (cis+trans- or cis *×*trans-acting) based on the results of the two significance tests. A gene is classified as conserved if neither the cis-acting effect nor the trans-acting effect are significantly different from zero. Cis+trans-acting and cis *×* trans-acting effects are classified if both the cis-acting and trans-acting effects are significant, and are then as-signed based on the direction of gene expression divergence. While assigning genes to discrete classification groups provides a useful summary for comparing cell types and tissues, it does not capture the full continuum of regulatory architecture. Hallgrímsdóttir et al., 2024 also introduced a proportion cis metric, which quantifies the relative contribution of cis-acting versus trans-acting regulation to differences in parental expression and can be analyzed alongside the coarser regulatory assignments.

To determine regulatory composition at the whole-tissue level, we also applied XgeneR to pseudobulked profiles of nine core tissues in each of the seven parental-F1 trios (Fig. 1D). Across all regulatory classes, we observed greater variability in regulatory proportions across tissues than across genotypes. Conserved regulation comprised 37-95% of genes across tissues and crosses, with the lowest proportion observed in the cortex and hippocampus and the highest in heart. Among non-conserved genes, cis-acting variation was the predominant class (72% average) followed by transacting variation (14% average), with the notable exception of B6J-PWK trios in gastrocnemius, where the proportion of trans-acting effects exceeded cis-acting. Cis *×*trans-acting and cis+trans-acting assignments were relatively rare, comprising an average of 9.1% and 4.0% of non-conserved regulatory effects.

To assess parent-of-origin effects on allele-specific expression, we identified candidate imprinted genes in pseudobulked whole-tissue samples by selecting loci exhibiting minimal parental divergence but strong allelic imbalance in F1 mice ( | parental log_2_ fold change| ≤ 1, | hybrid log_2_ fold change| ≥ 4, and |hybrid log_2_ fold change| *™* |parental log_2_ fold change| *≥* 3) (Fig. S3). Using this framework, we identified 83 candidate imprinted genes, including 37 long non-coding RNAs. Scatterplots comparing parental and hybrid log_2_ fold changes across tissues revealed a distinct subset of candidate imprinted genes separated from the broader distribution of regulatory effects by strong parentally biased expression patterns (Fig. S3A). Heatmap analysis demonstrated tissue- and strain-specific allelic bias among candidate imprinted genes, including known imprinted genes (Fig. S3B). Across all tissues and strains, paternally biased candidate imprinted genes were detected more frequently than maternally biased candidates (Fig. S3C,D), with the highest numbers observed in the diencephalon/pituitary gland and relatively few detected in liver, heart, and kidney. Canonical imprinted genes including *Airn, Rian*, and *Grb10* displayed robust parentally biased expression across multiple tissues and genetic backgrounds (Fig. S3E-G), consistent with previously described imprinting patterns.

While whole-tissue regulatory profiles aggregate diverse cell populations, single-cell resolution reveals distinct architectures based on tissue composition. We first examined the relationship between cell-type abundance and bulk-tissue concordance using hepatocytes, the dominant cell type in the liver at approximately 80% composition, and astrocytes, a relatively minor population at approximately 14% composition in the diencephalon (Fig. 1E-H). Hepatocytes showed high regulatory conservation, with 49% in CAST trios and 66% in WSB trios, and strong concordance with bulk liver results ( 82% sharing; Fig. S4A,B). In contrast, astrocytes displayed lower conservation (38–44%) and sharply diverged from bulk diencephalon profiles (25–29% sharing; Fig. S4C,D). This trend extended to other populations: liver cholangiocytes, a minor population, showed even less sharing with bulk liver (Fig. S4E,F), while glutamatergic neurons, an abundant brain cell type population, showed higher bulk concordance than astrocytes (Fig. S4G,H). Across all eight comparisons, regulatory patterns were more similar within a given cell type across strains than between different cell types, suggesting that cellular identity is a primary driver of regulatory architecture.

### Variation of cis-acting and trans-acting effects between brain regions in B6J-CAST trios

To explore how cis- and trans-acting effects vary across biological contexts, we focused on two functionally distinct brain regions: the cortex and hippocampus, which are essential to learning and memory, and the diencephalon, including the pituitary gland, which together serve as regulators of the nervous and endocrine systems. While these regions share several overlapping cell types, their divergent roles provide a tractable context for comparing regulatory architecture between shared cell types across anatomical regions. We focused on B6J-CAST parental-F1 trios, which represent the most genetically divergent mouse cross in our dataset. Additionally, the established behavioral and neurological differences between these strains provide a compelling biological context for comparing their regulatory landscapes^31,32^.

Across both regions, the regulatory assignment of majority genes in most cell types was classified as conserved. Notable exceptions were observed in several non-neuronal cell types, including microglia, mature oligodendrocytes, and endothelial cells across both regions; similar patterns were also observed in glutamatergic neurons of the cortex and hippocampus, and in astrocytes, oligodendrocyte precursor cells, and vascular leptomeningeal cells of the diencephalon, where non-conserved regulatory classes comprised a larger fraction of genes (Fig. 2A,B). Among genes with non-conserved regulation, cis-acting variation predominated in the cortex and hippocampus cell types and in most diencephalon cell types, with exceptions including microglial cells of both regions, and astrocytes, oligodendrocyte precursor cells, and endothelial cells of the diencephalon. This increased regulatory divergence in microglia is notable given their high degree of cellular plasticity and sensitivity to the local environment compared with other neuronal cell types^33^. Consistent with this pattern, diencephalon cell types contained a higher proportion of cis *×*trans (compensatory) regulatory assignments compared to cortex and hippocampus cell types. Importantly, in both tissues, differences in regulatory composition were not driven by the number of genes included in each analysis, as cell types with markedly different gene counts did not show a consistent trend in regulatory composition.

**Figure 2.**
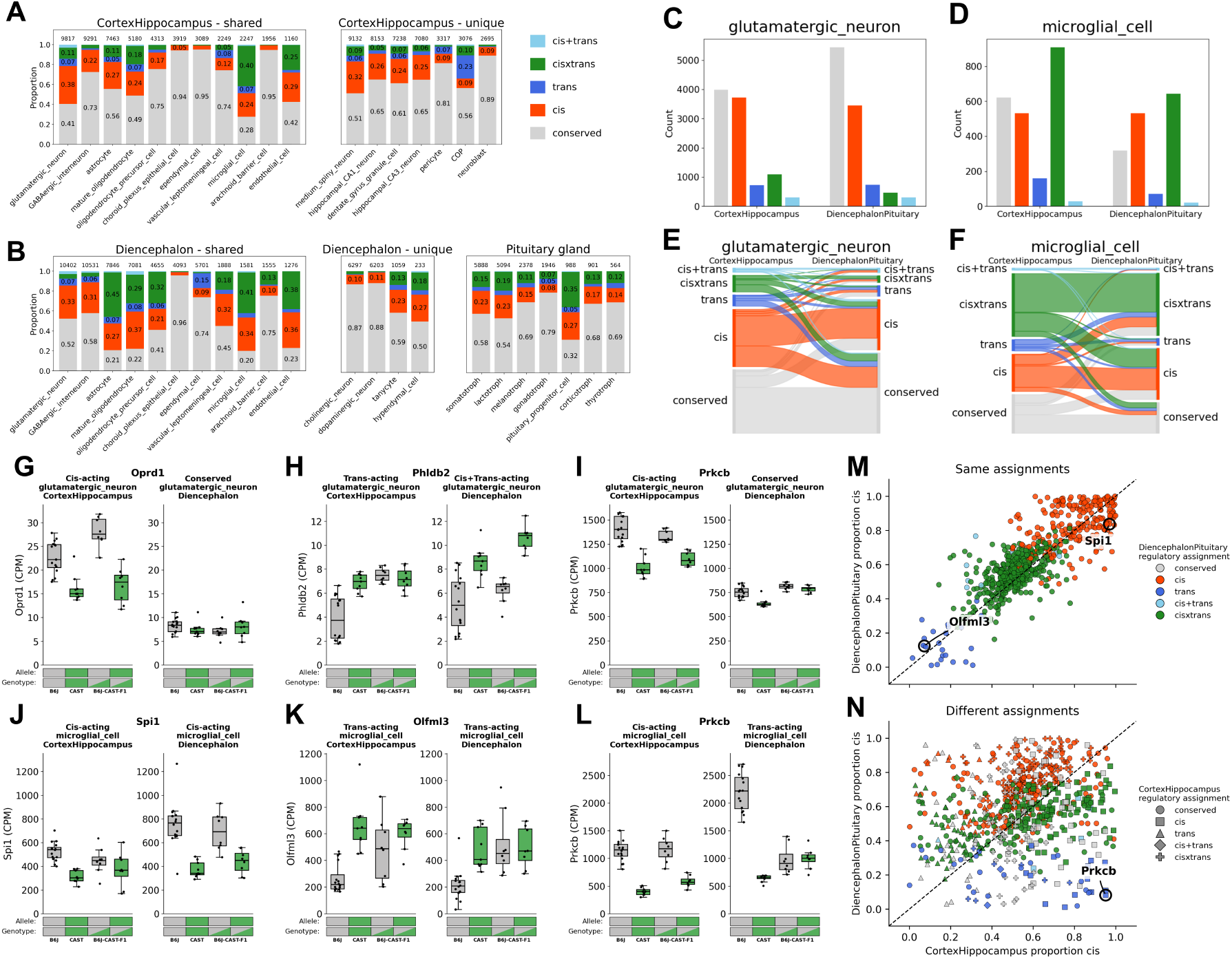
Cell-type resolved regulatory assignments in the cortex/hippocampus and diencephalon/pituitary gland. Proportion bar plots of genes in each regulatory category per cell type in the cortex/hippocampus (A) and the diencephalon/pituitary gland (B). Number of genes assigned to each regulatory category in the cortex/hippocampus and diencephalon for glutamatergic neurons (C) and microglia (D). Sankey diagrams showing the switches in regulatory assignments between the cortex/hippocampus (left) and diencephalon (right) for glutamatergic neurons (9,135 shared genes) (E) and microglia (1,558 shared genes) (F). Pseudobulked gene expression of B6J and CAST parental strains and allele-specific expression of B6J–CAST F1s, shown for glutamatergic neurons (G–I; Orpd1, Phldb2, Prkcb) and glutamatergic neurons (J–L; Spi1, Olfml3, Prkcb). Scatterplot of proportion cis values for microglial cells in each brain region for genes which share regulatory assignments (M) and different regulatory assignments (N).

We next focused on glutamatergic neurons and microglial cells, two well defined cell types, to examine regional differences in cis- and trans-acting regulatory architecture. Overall, both glutamatergic neurons and microglial cells contained similar numbers of genes with non-conserved regulatory assignments between tissues (Fig. 2C,D). Between the cortex/hippocampus and diencephalon, 9,135 genes in glutamatergic neurons and 1,558 genes in microglial cells were commonly detected across both regions. For both cell types, the majority of genes retained the same regulatory assignments between tissues (Fig. 2E,F). Among genes that differed in regulatory assignment between regions, transition frequency between orthogonal regulatory assignment (cisacting and trans-acting) was higher in glutamatergic neurons than microglial cells. Across tissues, in microglial cells, genes classified as cis- or trans-acting transitioned to cis*×* trans regulation 5.3-fold more frequently than they switched between cis and trans; in contrast, in glutamatergic neurons the opposite pattern was observed, with switches between cis-acting and trans-acting occurring 2-fold more frequently than transitions to cis×trans. Both microglial cells and glutamatergic neurons demonstrated similar concordance of regulatory assignments, sharing 54% of assignments between brain regions. These results indicate that regulatory assignments are largely conserved across brain regions, with some genes exhibiting tissue-specific transitions.

To illustrate how these regulatory assignments manifest at the level of individual genes, we examined representative examples of stable and context-dependent regulation in microglia and glutamatergic neurons. For example, the *δ*-opioid receptor *Oprd1* demonstrated cis-acting variation in glutamatergic neurons of the cortex/hippocampus but showed no differences in expression in glutamatergic neurons of the diencephalon (Fig. 2G). *Phldb2*, a gene required for projection stabilization, exhibited transitions between trans-acting regulation in the cortex/hippocampus and cis-acting regulation in the diencephalon (Fig. 2H). *Prkcb*, which encodes a signaling protein, was identified as cis-acting in the cortex/hippocampus and conserved in the diencephalon (Fig. 2I).

We next examined specific genes in microglial cells, such as the master regulator *Spi1*, which was classified as cis-acting in both brain regions, and the microglial marker gene *Olfml3*, which exhibited trans-acting variation in both regions, providing examples of shared modes of regulation between regions (Fig. 2J,K).^34^ In microglial cells, *Prkcb* was identified as being under cis-acting variation in the cortex/hippocampus and trans-acting variation in the diencephalon. Notably, *Prkcb* demonstrated consistent regulatory patterns between microglial cells and glutamatergic neurons in the cortex/hippocampus, and divergent patterns in the diencephalon (Fig. 2I,L).

To investigate changes in cis-acting contributions to gene expression differences, we examined proportion cis values for genes in microglial cells between brain regions. As expected, genes that shared regulatory assignments in microglial cells between the cortex/hippocampus and dien-cephalon, including *Spi1* and *Olfml3*, had similar proportion cis values (Fig. 2M). Genes with different regulatory assignments between brain regions clustered in the center of the proportion cis spectrum, consistent with an enrichment of regulatory switching between cis*×* trans- and cis- or trans-acting assignments. While *Prkcb* demonstrated divergent proportion cis values between the cortex/hippocampus and the diencephalon, most genes with differing regulatory assignments between regions exhibited moderate shifts in proportion cis values, consistent with the majority of transitions occurring between cis *×*trans-acting and cis- or transacting assignments (Fig. 2N). Together, these results demonstrate both shared and region-specific patterns of regulatory variation between the cortex/hippocampus and diencephalon, highlighting both cell-type- and tissue-specific regulatory differences.

### Broad regulatory variation across diverse cell types in B6J-CAST trios

To determine the distribution of regulatory patterns across the full cellular landscape, we next examined global patterns of cis-acting and trans-acting regulatory variation across all cell types in our dataset within B6J-CAST trios. We first examined pairwise relationships between regulatory class proportions across cell types (Fig. 3A). Because cis+trans additive regulation was consistently rare in all cell types, we focused this analysis on conserved, cis-acting, trans-acting, and cis *×* trans-acting regulatory classes. Across the 126 cell type–tissue combinations included in the analysis, regulatory composition varied substantially in a cell-type-specific manner (Fig. S5). The proportion of conserved regulation ranged from 20% in diencephalon microglia to 99% in heart adipocytes (Fig. 3A,B). The proportion of cis-acting genes ranged from 0% to 44% in adrenal endothelial cells.Cis*×* trans-acting regulation displayed a slightly wider range of variability, ranging from *<* 1% in heart macrophages to 52% in kidney endothelial cells. Trans-acting proportions were *<* 15% in all cell types except committed oligodendrocyte precursor cells of the cortex and hippocampus (23%) and liver cholangiocytes (28%). Cis+trans-acting variation showed the least variability overall, with the greatest proportion detected in liver cholangiocytes at 4% of genes. Together, these results reveal the extensive cell-type-specific variation in regulatory composition at single-cell resolution that is not apparent in analyses of bulk tissues.

**Figure 3.**
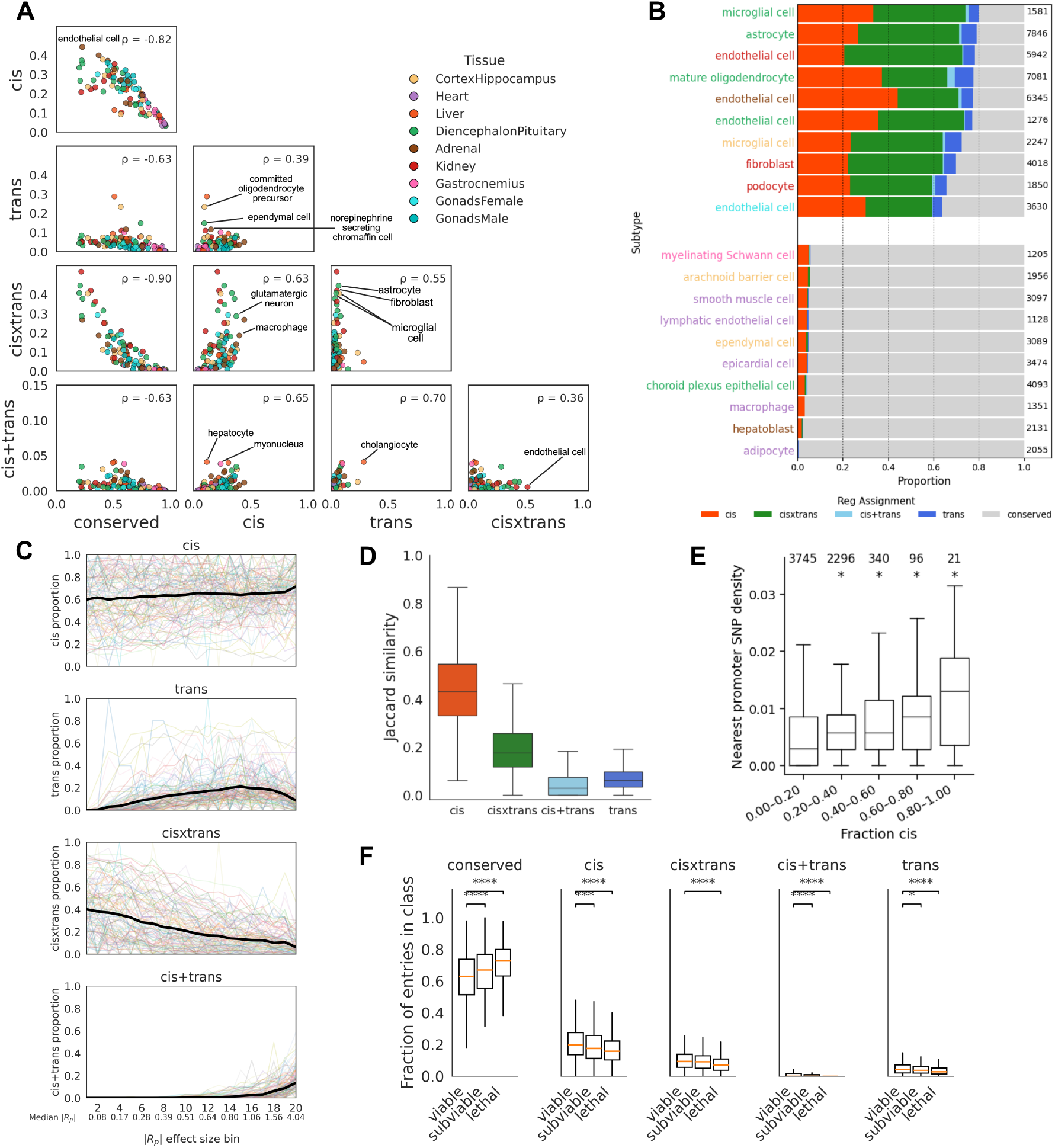
Regulatory variation across diverse cell types in B6J–CAST F1 trios. (A) Pairwise scatter plots showing proportion of regulatory assignments across nine tissues, encompassing 126 unique cell-type-tissue combinations. *ρ* indicates the Spearman correlation coefficient. (B) Stacked proportion bar plots of the top ten and bottom ten cell types by conserved proportion, with color of the cell-type text indicating its tissue of origin. (C) Proportions of regulatory assignments are plotted across 20 quantiles of absolute parental log_2_ fold change (*R*_*P*_ ). The black line indicates the global mean across all cell types. (D) Regulatory sharing between 5,506 cell-type pairs. (E) Distribution of promoter SNP density across levels of cis-regulatory conservation. Genes are binned by the fraction of cell types showing cis divergence. SNP density represents the proportion of SNPs within the proximal promoter region of the transcription start site (TSS). Mann–Whitney U test with Benjamini–Hochberg correction (* *q <* 0.05), with significance relative to the first bin. (F) Per-gene regulatory class fraction across 126 cell type–tissue combinations, grouped by IMPC knockout viability. Only genes observed in 33 combinations are shown. Mann–Whitney U test with Benjamini–Hochberg correction (* *q <* 0.05, ** *q <* 0.01, *** *q <* 0.001).

Because these proportions are compositional, such that all proportions sum to one, correlations between regulatory classes are not independent but can nonetheless reveal structured relationships across cell types. As expected, cis-acting and cis *×*trans-acting proportions varied inversely with conserved regulation. However, cis *×*trans proportions showed little direct association with either cis-acting or transacting proportions, indicating that cis *×* trans regulation does not simply scale with the prevalence of cis-acting or transacting effects (Fig. 3A). In contrast, trans-acting proportions showed little association with conserved regulation, remaining relatively stable across a wide range of non-conserved proportions, consistent with trans-acting regulation contributing a comparatively stable background component. Fur-thermore, cis+trans-acting variation was more strongly associated with an increase in trans-acting variation. Because more abundant cell types contain more nuclei, they may have greater power to detect regulatory differences. Consistent with this expectation, analysis of the relationship between the number of nuclei included in cell-type pseudobulk profiles and conserved proportions revealed a significant but weak correlation (Fig. S6).

In addition to the low levels of conservation observed in diencephalic microglia, we found that microglia of the cortex and hippocampus similarly showed low conservation (Fig. 3B). Three of the ten least conserved cell types were endothelial cells from different tissue groups. Together, these findings demonstrate that cell-type identity is a fundamental factor shaping how genetic variation impacts the regulatory landscape.

### Cell-type-specificity of regulatory variation

Having characterized how regulatory variation is distributed and related across cell types, we next examined properties not captured by compositional relationships. We first compared the magnitude of gene expression changes associated with each regulatory class by binning absolute parental log_2_ fold changes between B6J and CAST (Fig. 3C). Cis-acting variation contributed to the full range of parental gene expression divergence, with a modest increase in contribution to the largest magnitude differences, consistent with cis-regulatory variation contributing disproportionately to large heritable expression effect sizes.^35,36^ In contrast, cis*×* trans-acting variation preferentially contributed to small expression differences and progressively declined with increasing parental divergence, in line with antagonistic or compensatory regulatory interactions. Trans-acting variation contributed primarily to intermediate effect sizes, whereas cis+trans-acting variation, although relatively uncommon overall, became increasingly enriched among genes with the largest parental expression differences, indicating that reinforcing cis- and trans-acting effects are associated with the largest expression differences.

We next quantified cell-type specificity across regulatory classes by comparing regulatory assignments using Jaccard similarity between pairs of cell types sharing at least 50 non-conserved genes (Fig. 3D). Across 5,506 cell-type pairs, cis-acting variation was more frequently shared between cell types, with cis+trans- and trans-acting effects showing the highest degree of cell-type specificity. Trends were broadly consistent between individual cell-type pairs (Fig. S7). Notably, we observed inter-tissue variability in regulatory sharing between cell types, with cell types of the gastrocnemius exhibiting relatively large amounts of sharing of genes regulated by both cis-acting and trans-acting variation, a trend that was consistent across all parental–F1 trios (Fig. S8). These results demonstrate that trans-acting regulation is strongly cell-type-specific, likely reflecting differences in cell-type-specific regulatory environments, whereas cis-acting variation is encoded in local regulatory elements and is more broadly shared across cell types.

Having established that trans-acting and cis-and-trans-acting variation are more cell-type-specific than cis-acting variation, we next sought to identify properties underlying this difference. We hypothesized that genes classified as cis-acting across many cell types are driven by sequence vari-ation in promoter-like elements. To test this, we considered genes detected in at least 33 cell types (the median) and binned genes by the fraction of cell types in which they were classified as cis-acting. We then compared this fraction to SNP burden in the promoter region nearest the transcription start site (Fig. 3E). Genes more consistently classified as cis-acting across cell types exhibited a higher promoter SNP burden, suggesting that mutations in core promoter elements contribute to regulatory effects that are broadly shared across cellular contexts.

### Selective constraint shapes regulatory variation across cell types

Next, we examined the relationship between regulatory conservation and gene essentiality by assessing loss-of-function tolerance across regulatory classes^37^ (Fig. 3F). Of the genes included in our analysis, 7,088 had available knockout viability annotations. We restricted this set to genes detected in at least 33 cell types, and compared the fraction of cell types in which each gene was assigned to a given regulatory class with its knockout viability category (viable, sub-viable, or lethal). Genes classified as conserved were enriched among sub-viable and lethal categories, whereas genes assigned to non-conserved regulatory classes were more frequently observed among knockout-viable genes, with progressively lower representation in sub-viable and lethal categories. Notably, cis *×*trans-acting genes showed no significant difference between viable and sub-viable categories but were depleted among lethal genes, indicating that cis *×*trans regulation is tolerated in genes with partial constraint but is reduced in genes under the strongest selective pressure (Fig. S9). Together, these findings support a model in which genes central to development and core biological processes are subject to stronger selective constraint and therefore exhibit reduced mutational tolerance.

To further relate regulatory conservation to gene-level functional properties, we compared our results with cell-type specificity metrics from Swarna et al. (Fig. S10). *ζ* is a measure of how specifically a gene’s expression is restricted to one or a few cell types versus being broadly distributed across many cell types. Broadly expressed genes (low *ζ*) were enriched for conserved regulation, whereas genes with higher *ζ* showed increasing proportions of non-conserved regulation (Fig. S10A). We next classified genes as cell-type-specific using a block specificity score *>* 0.9 and restricted the analysis to cell types with at least 80 cell-type-specific genes. Cell-type-specific genes exhibited altered regulatory compositions relative to broadly expressed genes, including reduced conserved regulation in mature oligodendrocytes and increased cis+trans-acting regulation in several cell types, although these effects varied across regulatory classes and cell types, particularly in microglia (Fig. S10B,C).

### Cross-trio analysis reveals increased cis-acting variation in more divergent crosses

Having established the diversity of regulatory variation across cell types and tissues in B6J-CAST trios, we next examined how regulatory architecture varies across diverse genetic backgrounds. We first focused on microglia and glutamatergic neurons, two cell types analyzed extensively above, and compared regulatory composition across all seven parental-F1 trios (Fig. 4A,B). In glutamatergic neurons, the proportion of conserved genes ranged from 58% in 129S1 trios to 37% in PWK trios,with cis-acting variation consistently representing the largest non-conserved class (Fig. 4A). In microglial cells, conservation varied more widely, ranging from 68% in NZO and NOD trios to 28% in CAST trios (Fig. 4B). In contrast to conserved gene expression, both cis-acting and cis *×*transacting variation varied substantially across trios, with CAST trios exhibiting the highest proportion of cis *×*trans regulation. Across both cell types, trans-acting variation showed the least variability between genetic backgrounds.

**Figure 4.**
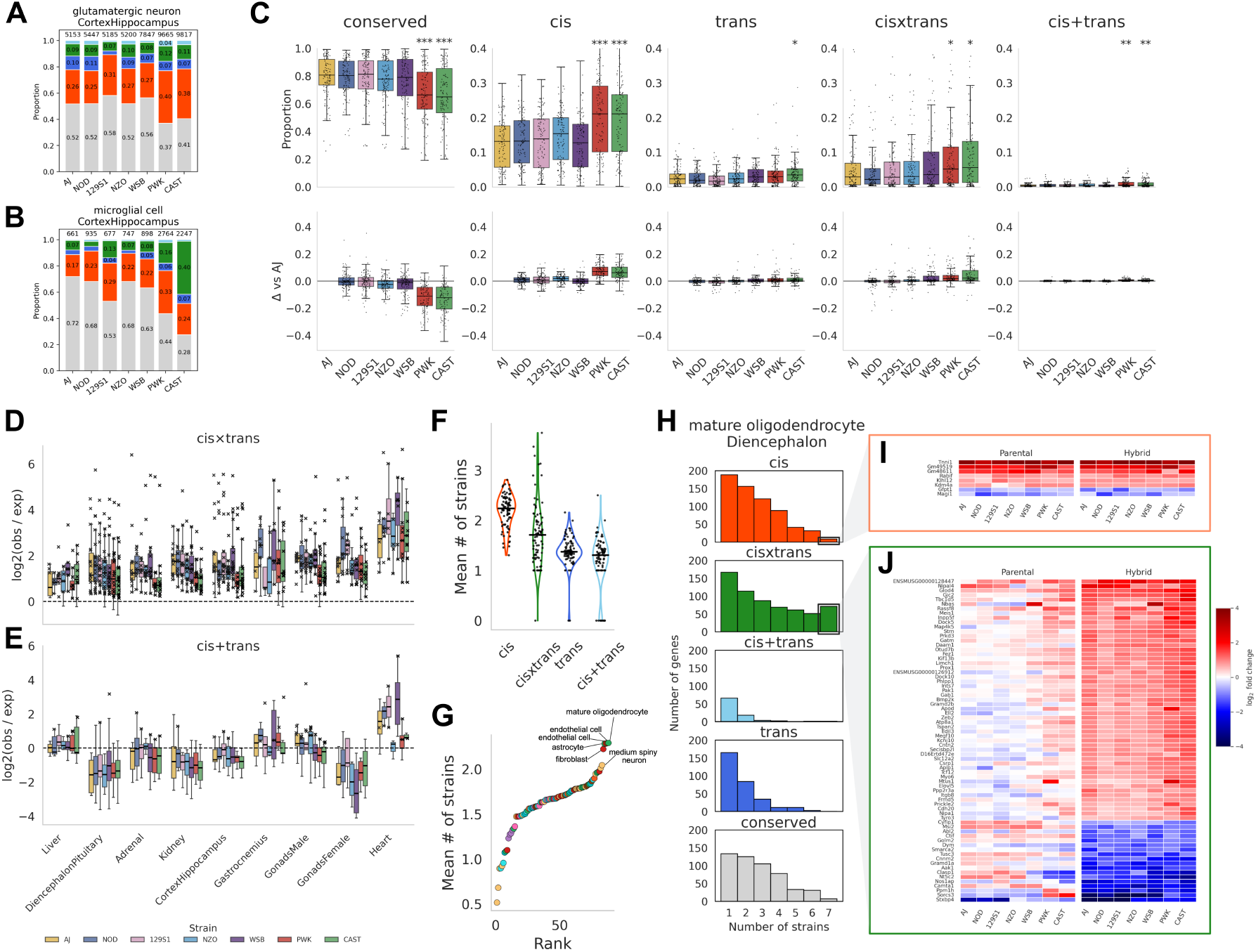
Landscape of cis- and trans-acting variation in seven parental–F1 trios. (A, B) Proportion bar plots showing the regulatory composition of glutamatergic neurons (A) and microglia (B) across seven parental–F1 trios. (C) Box plots (top) showing the proportion of genes within each regulatory class across all cell types for seven mouse strains, with significance relative to AJ (*p <* 0.01, Mann–Whitney U test). The corresponding divergence for each strain (bottom) is expressed as the *[INSERT METRIC]* in cell-type proportions relative to the AJ baseline. (D, E) Enrichment of observed cis trans-acting (D) and cis+trans-acting (E) variation relative to expected frequency. Box plots show the distribution across all cell types in each tissue. Crosses indicate individual cell types with significant enrichment (FDR *<* 0.05, Fisher’s exact test). (F) Mean number of strains in which a gene shares a regulatory assignment for each cell type, shown separately for cis-, cis*×*trans-, and trans-acting assignments. (G) Overall mean sharing of regulatory assignments across all cell types. (H) Number of parental–F1 trios in which a gene shares a regulatory assignment for mature oligodendrocytes of the diencephalon. (I, J) Heatmaps of parental and hybrid log_2_ fold changes for genes with shared regulatory assignments across all seven parental–F1 trios, including 8 genes with shared cis-acting (I) and 71 genes with shared cis*×*trans-acting (J) assignments.

To assess how genetic background influences regulatory composition globally across cell types, we compared cell-type-level proportions of each regulatory class across the seven parental–F1 trios, using AJ trios as a reference because they exhibit the least genetic divergence from B6J (Fig. 4C). Among *domesticus* subspecies trios, we observed no significant differences in the proportions of any regulatory class relative to AJ. In contrast, PWK and CAST trios exhibited significantly reduced conserved regulation, with average decreases of 11.0% and 12.1%, respectively. The observed decrease in conserved gene expression in PWK and CAST was largely explained by an increase in cis-acting variation, with smaller contributions from increased cis*×* trans- and cis+trans-acting variation. These results suggest that variants acting in cis preferentially drive gene expression divergence with increasing genetic distance.

### Regulatory class sharing between crosses suggests shared regulatory mechanisms

To determine whether combined cis- and trans-acting regulatory variation arises beyond expectations from independent cis- and trans-acting effects, we calculated the ratio of observed to expected genes classified as either cis *×*trans- or cis+trans-acting for each cell type across tissues and strains (Fig. 4D,E). Across all seven crosses, cis *×*trans-acting variation occurred more frequently than expected based on the individual frequencies of cis and trans effects, with enrichment varying primarily by tissue rather than by strain. Heart and gonadal tissues showed the strongest enrichment, whereas liver and skeletal muscle exhibited comparatively weaker signals. In contrast, cis+trans-acting variation generally occurred at or below expected levels across most tissues and cell types, and comparatively few cell types showed significant enrichment. The contrasting patterns observed for cis *×*trans and cis+trans variation may reflect differing evolutionary constraints on compensatory versus reinforcing regulatory effects.

Our dataset spans seven genetically distinct parental– F1 crosses, enabling direct comparison of regulatory sharing across genetic backgrounds. For each regulatory class, we calculated the average number of strains in which a gene shared the same regulatory assignment within a given cell type (Fig. 4F). Genes classified as cis-acting were more frequently shared across strains than genes classified as cis-and- trans- or trans-acting. Across cell types, mature oligodendrocytes of the diencephalon showed the highest overall degree of regulatory sharing, followed by endothelial cells of the kidney and adrenal gland and astrocytes of the diencephalon (Fig. 4G). We identified 666 genes that shared a non-conserved regulatory assignment among the seven parental– F1 trios in at least one cell type. Overall, we found moderate levels of regulatory sharing within cell types across the different crosses.

Because mature oligodendrocytes exhibited the highest overall degree of regulatory sharing, we used this cell type as a representative case to examine the distribution of genes shared across strains (Fig. 4H). As expected, most genes shared regulatory assignments across more than one trio. We identified 8 cis-acting genes and 71 cis *×*trans-acting genes that retained the same regulatory assignment across all seven trios. These genes are notable in that they suggest shared regulatory variants among the non-B6J strains, with B6J representing the divergent allele. Consistent with this interpretation, the 8 cis-acting genes shared across all seven trios showed shared direction of effect in all crosses (Fig. 4I). In contrast, the 71 cis*×*trans-acting genes exhibited small parental log_2_ fold changes and consistent hybrid effect directionality, congruent with expectations of compensatory regulatory interactions (Fig. 4J). These shared regulatory effects across strains are suggestive of conserved regulatory variation in different genetic backgrounds.

## Discussion

A major goal of genomics is to characterize the genetic architecture that governs the effects of genetic variation on gene expression across tissue and cell-type contexts. While we previously established widespread cell-type-specific variation in gene expression between diverse mouse strains^25^, whether these changes were driven primarily by cis-acting or trans-acting genetic variants remained unresolved. Our classification of gene expression differences into cis-acting and trans-acting components across eight tissue groups and seven mouse parental-hybrid trios reveals that cis-acting variation is the predominant driver of gene expression divergence among cell types, whereas trans-acting effects are more restricted and more tightly coupled to cellular identity.

Resolving gene-expression architecture at cell-type resolution reveals substantial heterogeneity masked by bulk-tissue assays. While bulk-tissue analyses^5^, including our own, have highlighted cis-acting variants as the main drivers of expression divergence, cell-type-resolved analysis uncovers a much more heterogeneous regulatory landscape. For example, while cis-acting effects are frequently the largest non-conserved mode of regulation, certain cell types exhibit notably higher proportions of trans-acting or cis *×*transacting variation. Furthermore, our observation that cis-acting variation expands with genetic distance while transacting effects remain relatively stable is consistent with recent re-evaluations of how regulatory architecture diverges with evolutionary distance^5^. Ultimately, these results emphasize that a complete characterization of genetic diversity requires moving beyond aggregate bulk-tissue or whole-organism measures to account for the diverse regulatory landscape present across individual cell types and distinct genetic backgrounds.

We find that genes with cis-acting effects are more likely to be detected in multiple cell types than are trans-acting or cis-and-trans-acting interactions. This observation underscores a difference in regulatory logic: cis-acting variants are encoded locally in the genome, potentially modifying cis-regulatory elements and altering the expression of nearby genes, whereas the action of trans-acting variants depends on a cell’s regulatory environment. For example, a genetic variant that changes the expression of a transcription factor can have downstream propagated effects on target gene expression only if that transcription factor is transcribed and translated in a particular cellular context^39^. By contrast, a mutation that disrupts a core binding motif within a gene’s promoter may exert a constitutive effect regardless of cellular context. This observation of cell-type specificity is consistent with the omnigenic model, which proposes that gene expression is shaped by networks of transacting peripheral genes, with the influence of these networks filtered through cell-type-specific regulatory environments^2^. More broadly, these results highlight that the effects of genetic variation are inherently context-dependent. To fully understand genetic diversity, we must therefore consider the range of factors that define the Gene *×*Environment interactions to include intrinsic factors such as cell type, tissue context, and anatomical region, along with extrinsic factors such as circadian rhythm, age, and diet. In other words, phenotype is a product of Gene*×*(cell type*×*tissue*×*anatomical region*×*circadian rhythm*×*age*×*diet*×···* ).

We observed a distinct relationship between gene essentiality and the mode of regulatory divergence. For genes classified with cis-, trans-, or cis+trans-acting effects, there is a progressive decrease in the frequency of these effects as a gene becomes more essential, transitioning from viable to sub-viable and ultimately to lethal. Notably, cis*×*trans-acting compensatory effects were significantly depleted among lethal genes but maintained similar frequencies between sub-viable and viable categories. This suggests that while viable and sub-viable genes possess the functional flexibility to accumulate compensating regulatory variants, lethal genes are subject to intense stabilizing selection. Because essential genes have a smaller tolerance for expression changes, they likely cannot survive the deleterious intermediate states required to eventually accumulate a second, compensatory mutation. These results imply that once a gene falls below a strict threshold of essentiality, it gains the functional slack necessary to acquire subsequent compensatory variants. Furthermore, the breadth of conservation observed across our dataset may be further explained by pleiotropic constraint with genes that are expressed across a wide array of cell types are more likely to exhibit conserved expression than those with cell-type-specific roles, suggesting that multi-contextual requirements limit permissible the space for regulatory evolution.

It is important to consider that C57BL/6J (B6J), despite its status as the standard laboratory reference strain^40^, has its own private set of variants that are not present in most mice. For example, B6J has been shown to contain a homozygous deletion in the *Nnt* gene, impacting the ability of B6J mice to metabolize exogenous peroxide^41^. Furthermore, B6J mice and other commonly used laboratory mouse strains are melatonin deficient due to two recessive mutations in *Hiomt* and *Aanat*^42–44^. Across seven hybrid trios, we identified 666 genes that were assigned a non-conserved regulatory class in at least one cell type, indicating that these effects are shared across trios and suggesting that B6J may be an evolutionary outlier at a subset of loci. For example, in mature oligodendrocytes, we identified sets of cis-acting and cis*×* transacting genes for which all seven non-B6J strains exhibit concordant regulatory patterns that diverge from the B6J allele. These findings underscore the necessity of incorporating diverse mouse subspecies into studies of gene regulation, as observations made in a single mouse strain may reflect lineage-specific features rather than generalizable principles.

While expression quantitative trait locus (eQTL) mapping remains the standard approach for identifying genomic loci associated with phenotypic variation in humans and other natural populations, the framework applied here provides a mechanistically complementary approach^45^. Unlike traditional eQTL studies, which require large cohorts and often have limited power to detect trans-acting or compensatory regulatory effects^46^, allele-specific analysis in parental-hybrid crosses enables direct dissection of cis- and trans-regulatory contributions^12^. Historically, eQTL studies have primarily mapped regulatory variation in bulk tissues^47,48^, with more recent extensions to single-cell resolution largely restricted to accessible human samples such as blood^49–51^. In contrast, our results demonstrate that resolving regulatory architecture at the level of individual cell types across diverse tissues is essential for capturing the full spectrum of heritable regulatory variation. Together, these findings underscore the need for expanded, cell-type-resolved mouse eQTL resources to fully characterize the genetic architecture underlying gene-expression variability.

## Data and Code availability

- IGVF measurement sets for raw fastqs and uniformly processed AnnData h5ad files are listed in Supp. Table 2.
- Notebooks for figure generation, cell type annotation, and data processing are accessible at: github.com/mortazavilab/cistrans_manuscript
- A containerized, interactive data viewer can be downloaded from: github.com/mortazavilab/mousaic

## Acknowledgements

We thank the UCI Transgenic Mouse Facility for housing the mice and use of their facilities and UCI GRTH for sequencing the libraries. The TMF and GRTH are shared resources of the Chao Family Comprehensive Cancer Center, supported in part by the National Cancer Institute of the National Institutes of Health under award number P30CA062203. A.M. and B.J.W. were supported by UM1HG012077.

## Author contributions

A.M., I.B.H, L.P., and B.J.W. designed the project. S.K. acquired mice and oversaw their husbandry. G.F. and H.Y.L. dissected tissues under guidance from B.A.W., G.R.M., and S.K. M.D., P.M., E.T., N.F., N.M., and R.M. helped with sample collection as well as performed nuclei isolation, barcoding, and library preparation. D.T. uploaded data and metadata to the IGVF portal. M.C., N.S., and E.R. provided input and code for data analysis. R.W. performed data analysis, generated figures, and wrote and edited the manuscript with significant input from A.M.

## Methods

### Mice and tissue collection

Mice were obtained from The Jackson Laboratory (Bar Harbor, ME) and housed at the UCI vivarium under controlled conditions. All animal procedures were approved by the UCI Institutional Animal Care and Use Committee. Metadata for each animal and tissue, including mouse ID, sex, date of birth and euthanasia, time of euthanasia, dissector ID, body and tissue weights, and estrus stage, and zeitgeber time (ZT), defined as the number of hours since lights on (ZT0) are detailed in Supplementary Table 1. Animals were housed in individually ventilated cages (Blue Line, Technoplast, Seaford, DE) containing corncob bedding (Envigo 7092BK 1/8” Teklad, Placentia, CA) and two 2” square cotton nestlets (Ancare, Bellmore, NY) plus a LifeSpan multi-level environmental enrichment platform. Tap water and food (Envigo Mouse 2027; LabDiet, St. Louis, MO) were provided ad libitum. Cages were changed every 2 weeks with a maximum of 5 adult animals per cage. Room temperature was maintained at 72 *±* 2°F, with ambient room humidity (average 40–60% RH, range 10–70%). Light cycle was 12h light / 12h dark, lights on at 06:30h and off at 18:30h.

Tissue collection was performed as described in Rebboah et al., 2025. Briefly, mice were euthanized via isoflurane anesthesia followed by decapitation. Tissues, including brain regions, trunk organs, and limb muscles were dissected in parallel by expert personnel. All tissues were flash-frozen in liquid nitrogen and stored at -135°C.

### Nuclei isolation and single-nucleus RNA-seq experiments

Nuclei isolation and single-nucleus RNA-sequencing experiments were carried out as described in Rebboah et al., 2025. Briefly, flash frozen tissues from male and female B6J and F1 mice were processed in groups of eight replicates (four male and four female) for each tissue. Tissues were mechanically dissociated, followed by filtering, resuspension, and DAPI staining for nuclei counting. Nuclei fixation was performed using Parse Biosciences’ Nuclei Fixation Kit v2 according to the manufacturer’s protocol. 4 million nuclei were fixed for each sample, except for smaller tissues including female gonads and adrenal glands in which 1 million nuclei were fixed instead. Nuclei were stored at -80°C following final counting.

Single-nucleus RNA-sequencing libraries were generated using eight Parse Biosciences’ Evercode WT Mega Kits (v2). Unlike the plate layout used in Rebboah et al., 2025, each of the eight barcoding plates were prepared and loaded with 64 replicates for a single tissue for a total of 512 samples. The first 64 wells of the plate were loaded with individual replicates, with the remaining 32 wells containing multiplexed pairs of the same 64 samples. Libraries were built per the manufacturer’s protocol and as described in detail in Rebboah et al., 2025, producing 15 subpools of 67,000 nominal nuclei per plate for all tissues with the exception of heart. Subpools were sequenced together with a single run of the Illumina NovaSeq X Plus, with an average depth of 25 billion reads per tissue (25,000 reads per nuclei). We recovered fewer than expected heart nuclei. Heart subpools were instead sequenced with an Illumina Nextseq 2000 to a depth of 1.8 billion reads.

### Data preprocessing

Single-genome reference mapping data were retrieved from the IGVF portal. snRNA-seq data were processed via the IGVF single-cell uniform mapping pipeline, utilizing the GRCm39 reference genome (IGVFFI9282QLXO) and GENCODE M36 annotations (IGVFFI4777RDZK). FASTQ files were mapped using the kallisto bustools suite^52–54^.

### Single genome mapping QC and cell type annotation

Post-mapping processing and QC was performed as previously described in Rebboah et al., 2025. Briefly, CellBender^27^ was used to remove ambient RNA and background noise. Scrublet was run on uncorrected counts to identify and exclude potential doublets^55^. Genetic demultiplexing on multiplexed wells was performed by Klue^56^. Tissue-level single-nuclei datasets were generated and subjected to quality control. Nuclei were included if they met the following criteria: *>* 500 and *<* 150,000 UMIs, *>* 250 expressed genes ( ≥1 UMI), *<* 1% mitochondrial gene expression, and *<* 0.25 doublet scores.

Downstream analysis was performed using Scanpy (v1.11.4)^57^. CellBender-corrected counts were normalized and filtered for highly variable genes prior to dimensionality reduction and Leiden clustering. Following clustering, nuclei were annotated according to expression of a curated marker gene atlas, utilizing Cell Ontology (CL) IDs for defined cell types^58^. We annotated 92 cell types and states across the eight major tissue groups included in this dataset.

### Allele specific mapping

To quantify allele-specific expression, we utilized a mouse strain-specific Variant Call Format (VCF) file (https://ftp.ebi.ac.uk/pub/databases/mousegenomes/REL-2112-v8-SNPs_Indels/mgp_REL2021_snps.vcf.gz)^28^ containing variant data for the seven non-B6J founder strains. Using g2gtools (https://github.com/churchill-lab/g2gtools), these variants were incorporated into the GRCm39 genome to generate a unique, strain-specific reference genome (FASTA) and annotation (GTF) file for each of the seven strains. To enable allele-specific quantification with kallisto, we then generated concatenated references by merging the standard GRCm39 files with each of the seven strain-specific reference pairs.

FASTQ files representing 252 subpools from both founder and F1 mice were acquired from the IGVF portal. To eliminate mapping biases, both homozygous parental strains and heterozygous F1 genotypes were processed identically. Reads from each subpool were aligned independently to all seven concatenated allele-specific reference genomes to generate allele-specific count mappings using kallisto bustools. During downstream analysis of the founder strains, only reads mapping to the expected reference allele were retained for analysis.

### Allele specific mapping QC

Following alignment, the resulting .h5ad files from each subpool were processed using Cell-Bender to computationally remove ambient RNA. To assign metadata to these allele-specific mappings, nuclei strain identities (for multiplexed wells) and cell type annotations were lifted over directly from the single-genome mapping results detailed previously. Strict quality control was maintained by retaining only those nuclei that had successfully passed QC filters in the single-genome analysis. Because all downstream quantitative analyses were designed for pseudobulk profiles, specific UMI count thresholds were not enforced on the allele-specific mappings at the single-nucleus level.

The datasets were then filtered by their respective strains and tissues to generate final tissue-level AnnData objects. For the founder datasets, this included retaining exclusively the allele-specific mappings for B6J and the corresponding non-B6J reference strain. For the F1 datasets, data was strictly filtered to contain only mappings for B6J and the relevant F1 strain. Finally, the allele-specific count results within these filtered AnnData objects were pseudobulked using decoupler^59^ to generate robust, cell type-level pseudobulk profiles for downstream analysis.

### Assignment of regulatory variation

To quantify the relative contributions of cis- and trans-acting genetic variation to gene expression divergence, we applied XgeneR (v0.2.0)^30^. This approach utilizes a generalized linear model applied to allele-specific expression (ASE) in F1 hybrids and parental expression estimates.

We ran XgeneR independently for each cell type within each of the seven F1 crosses in our dataset. To avoid low-count artifacts, we restricted analyses to genes with ≥5 counts per million in a cell type using single-genome mapping, and with a mean of ≥10 allele-specific counts across both parental and hybrid expression profiles. Genes located on the X and Y chromosomes were excluded due to monoallelic expression. Significant regulatory attributions to parental expression differences (cis-, trans-, cis×trans-, and cis+trans-acting) were identified using an adjusted p-value threshold *<* 0.05. Genes above the p-value threshold were assigned as conserved. Additionally, XgeneR calculates both parental and hybrid log_2_ fold-change, and quantifies the proportion cis to determine fractional contribution of cis-acting genetic effects on total expression divergence.

### Promoter SNP density analysis

CAST/EiJ variants were intersected with promoter-like signatures (PLS) from the ENCODE mm39 cCRE catalogue (v4)^60^ using PyRanges^61^ to compute a per-PLS SNP density (SNPs per bp). Each gene’s transcription start site was mapped to its nearest PLS, and the corresponding SNP density was assigned to the gene. Per-gene cis-regulatory assignments across 126 cell type-tissue combinations were aggregated to compute a fraction-cis score (proportion of combinations in which the gene was classified as cis-regulated in CAST F1 crosses). Genes observed in fewer than 33 cell type-tissue combinations (median) were excluded. Promoter SNP density was then compared against fraction-cis across genes.

### Regulatory classification and loss-of-function viability

Knockout viability data were obtained from the International Mouse Phenotyping Consortium (IMPC)^37^: https://ftp.ebi.ac.uk/pub/databases/impc/all-data-releases/latest/results/viability.csv.gz. For genes with multiple knockout results, the most severe viability outcome was retained (lethal *>* subviable *>* viable). Per-gene regulatory classifications were aggregated across 126 cell type-tissue combinations to compute the fraction of combinations in which each gene was assigned to each regulatory class. Genes observed in fewer than 33 cell type-tissue combinations were excluded. Differences in regulatory class fraction across viability groups were assessed using two-sided Mann-Whitney U tests comparing viable versus subviable and viable versus lethal genes within each regulatory class, with p-values corrected for multiple comparisons using the Benjamini–Hochberg procedure.

### Regulatory enrichment

Cis×trans-acting variation enrichment was determined by calculating the expected number of interactions under independence (expected = (number of genes with a cis-acting effects + all genes with a trans-acting) / all genes including conserved) and comparing this with the observed number of genes identified as being influenced by cis×trans-acting variation.

### Interactive data viewer

An interactive web application was developed in Python 3.11 using Streamlit for visualization and exploratory analysis of cis-acting and trans-acting regulatory results and allele-specific gene expression. Gene expression data were processed using AnnData with normalization to counts per million. Interactive plots were generated with Plotly; static figures with Matplotlib. The application was containerized using Docker.

## Supplementary Tables

- **Table S1. Sample collection metadata**.
- **Table S2. IGVF file accessions**.

## Supplementary Figures

**Figure S1.**
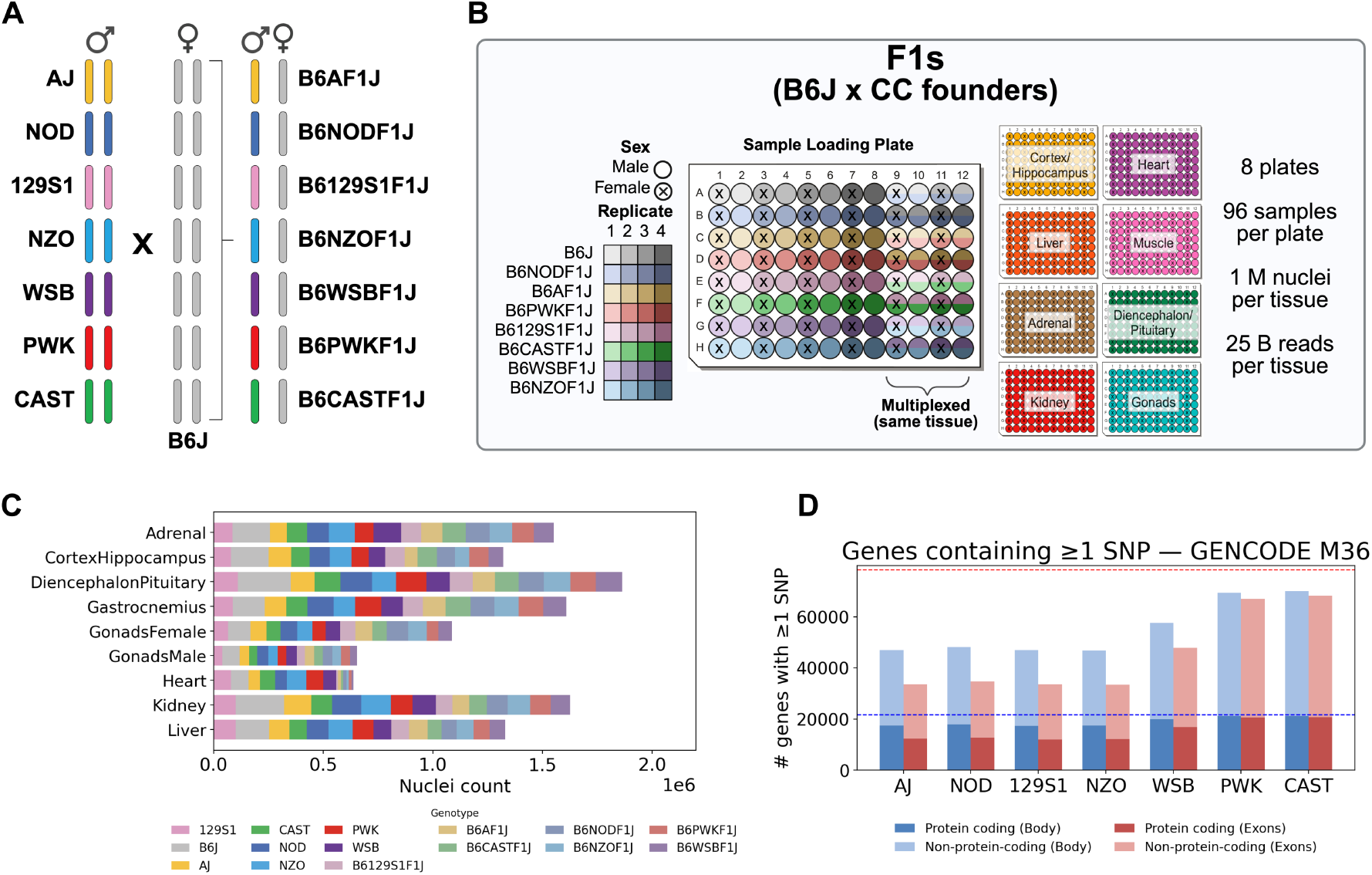
Experimental design. (A) Schematic representation of F1 crosses. (B) Design of sample barcoding plate for F1 samples. (C) Total number of nuclei passing quality control (QC) filters across nine tissues: Adrenal (n = 1,552,629), Cortex/Hippocampus (n = 1,321,104), Diencephalon/Pituitary (n = 1,863,454), Gastrocnemius (n = 1,608,883), Female Gonads (n = 1,087,721), Male Gonads (n = 654,238), Heart (n = 637,460), Kidney (n = 1,625,653), and Liver (n = 1,328,965). (D) Number of protein coding genes and non-protein coding genes which contain a SNP in exons or the full gene body. Horizontal lines indicate the total number of protein coding genes (blue; 21,748) and total annotated genes (red; 78,298) in GENCODE M36.

**Figure S2.**
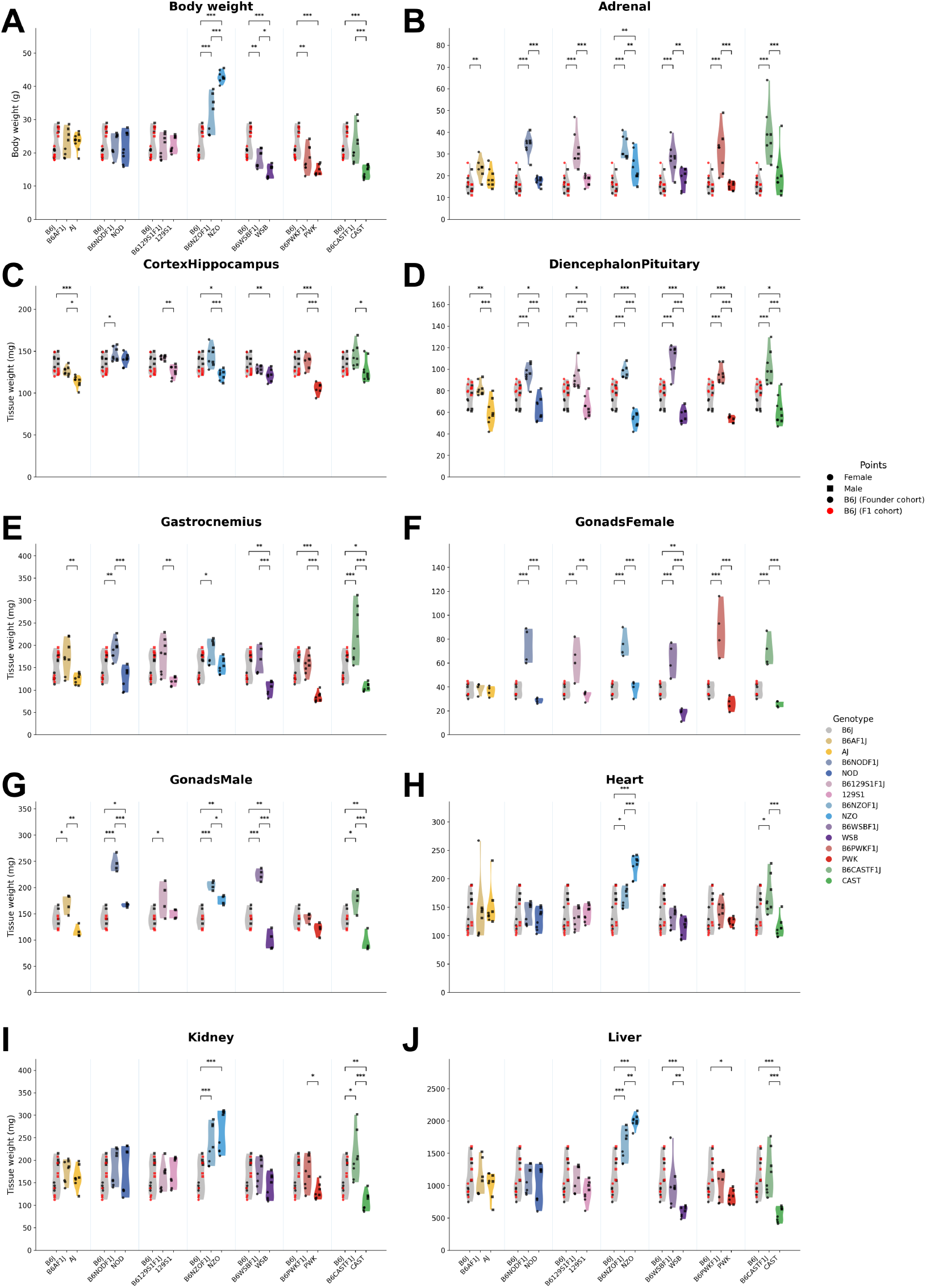
Body and tissue weight of the eight parental mouse strains and seven F1 hybrids. (A) Body weight, (B) Adrenal, (C) Cortex/hippocampus, (D) Diencephalon/pituitary gland, (E) Gastrocnemius, (F) Female gonads, (G) Male Gonads, (H) Heart, (I) Kidney, (J) Liver. Kidney, female gonad, and gastrocnemius weights for founder parental genotypes were divided by two, as left and right tissues were weighed together in the founder dataset, whereas F1 samples contained measurements from a single tissue. Pairwise differences between each genotype were assessed by one-way ANOVA followed by Tukey’s HSD post-hoc test. (* p < 0.05, ** p < 0.01, *** p < 0.001).

**Figure S3.**
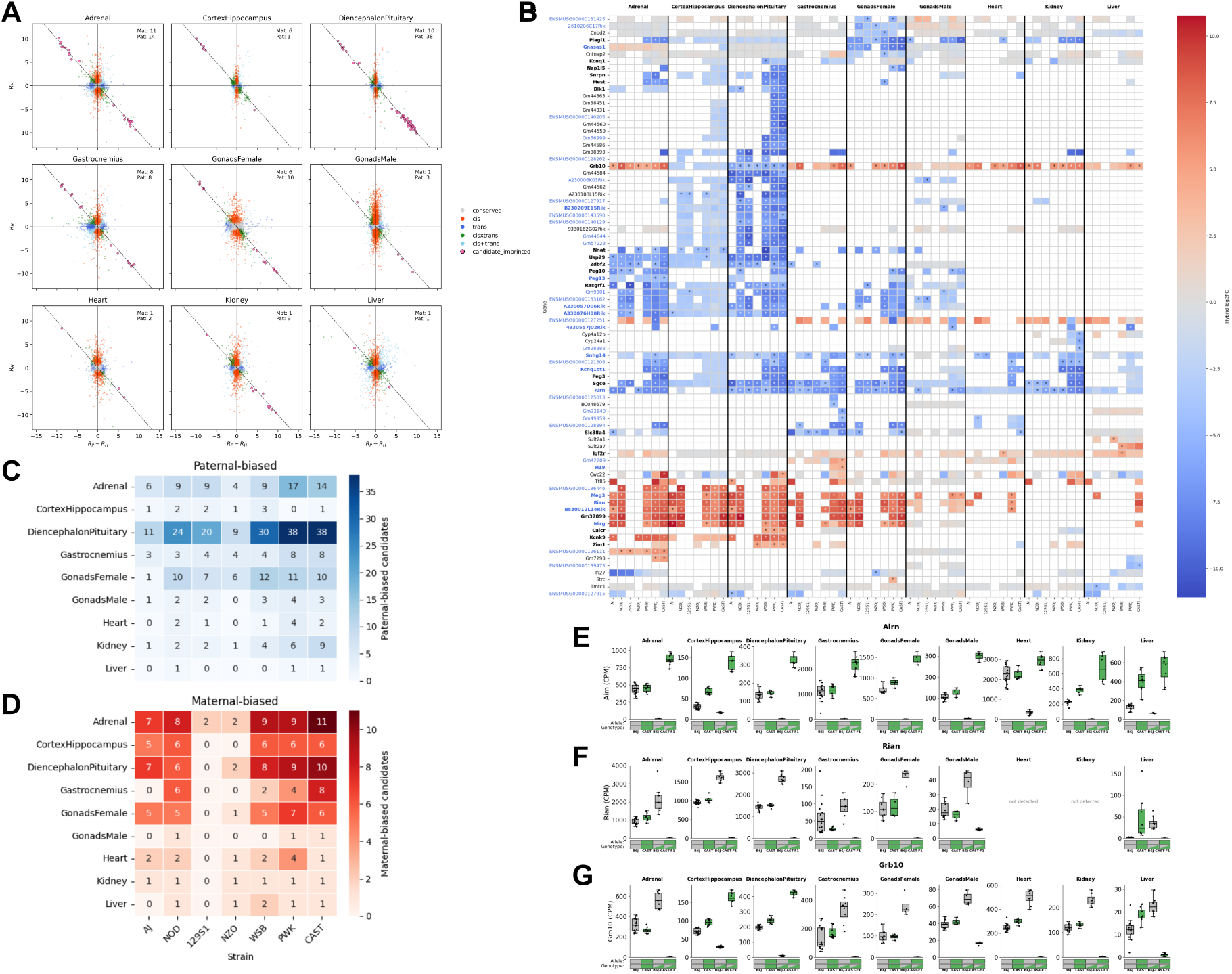
Identification of imprinting candidate genes. (A) Scatterplots comparing parental log_2_ fold changes (*R*_*P*_ ) and hybrid allelic log_2_ fold changes (*R*_*H*_ ) in CAST/EiJ crosses across nine tissues. Candidate imprinted genes, defined as genes with low parental divergence but strong hybrid allelic imbalance, are highlighted in pink. Numbers indicate the number of maternally (Mat) and paternally (Pat) biased candidates per tissue. (B) Heatmap of hybrid allelic imbalance (*R*_*H*_ ) for candidate imprinted genes across tissues and strains. Asterisks indicate tissue–strain combinations meeting candidate imprinting criteria. Known imprinted genes are shown in bold, and long non-coding RNAs are shown in blue. (C, D) Number of paternally biased (C) and maternally biased (D) candidate imprinted genes detected across tissues and strains. (E–G) Allele-resolved expression of representative candidate imprinted genes across tissues, including *Airn* (E), *Rian* (F), and *Grb10* (G).

**Figure S4.**
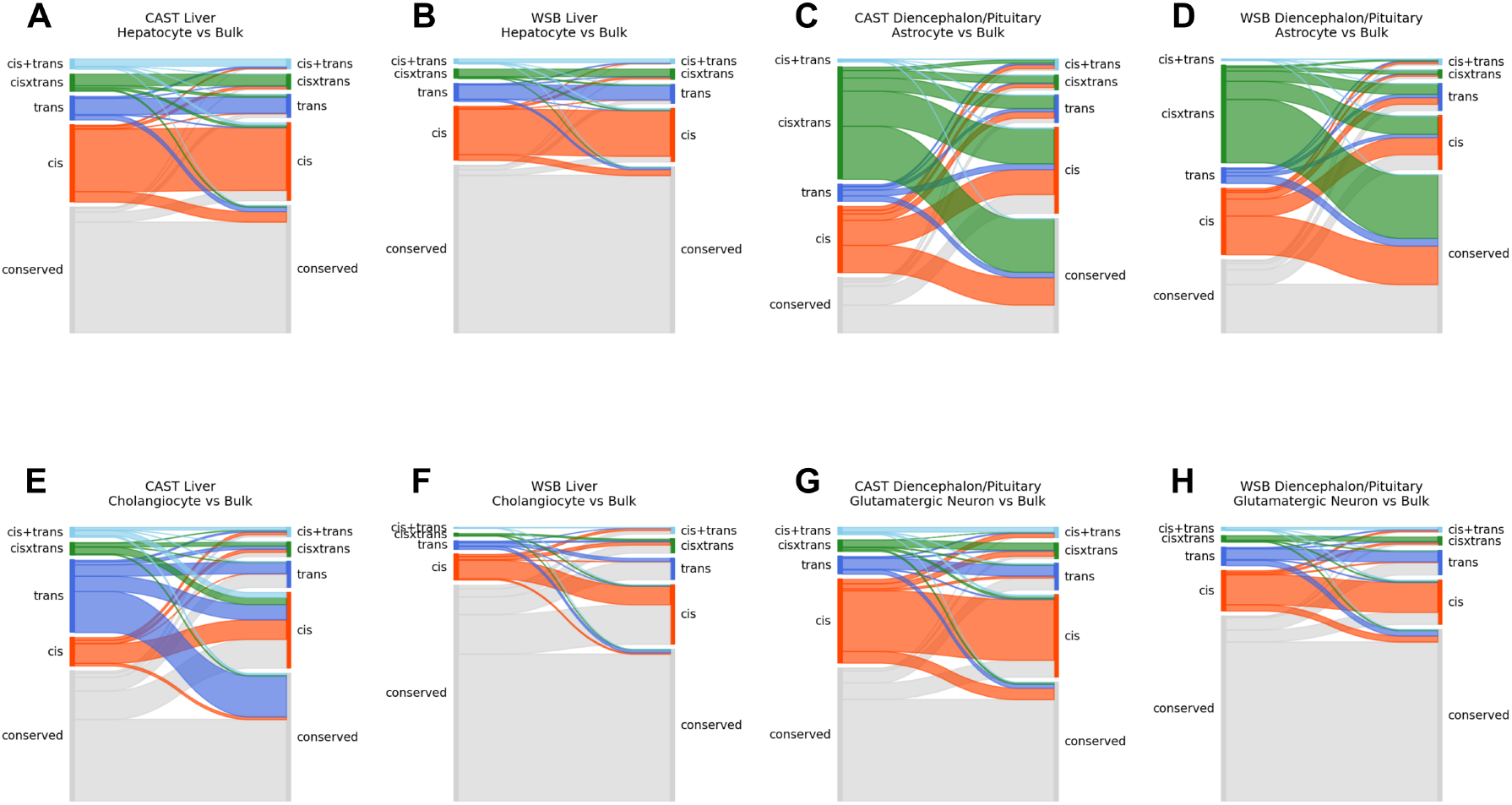
Regulatory assignment sharing between bulk tissue and cell types. (A) Hepatocytes (left) and bulk liver (right) in CAST trios, (B) Hepatocytes and bulk liver in WSB trios, (C) Astrocytes and bulk diencephalon/pituitary in CAST trios, (D) Astrocytes and bulk diencephalon/pituitary in WSB trios, (E) Cholangiocytes and bulk liver in CAST trios, (F) Cholangiocytes and bulk liver in WSB trios, (G) Glutamatergic neurons and bulk diencephalon/pituitary in CAST trios, (H) Glutamatergic neurons and bulk diencephalon/pituitary in WSB trios. See also figure 1D-H.

**Figure S5.**
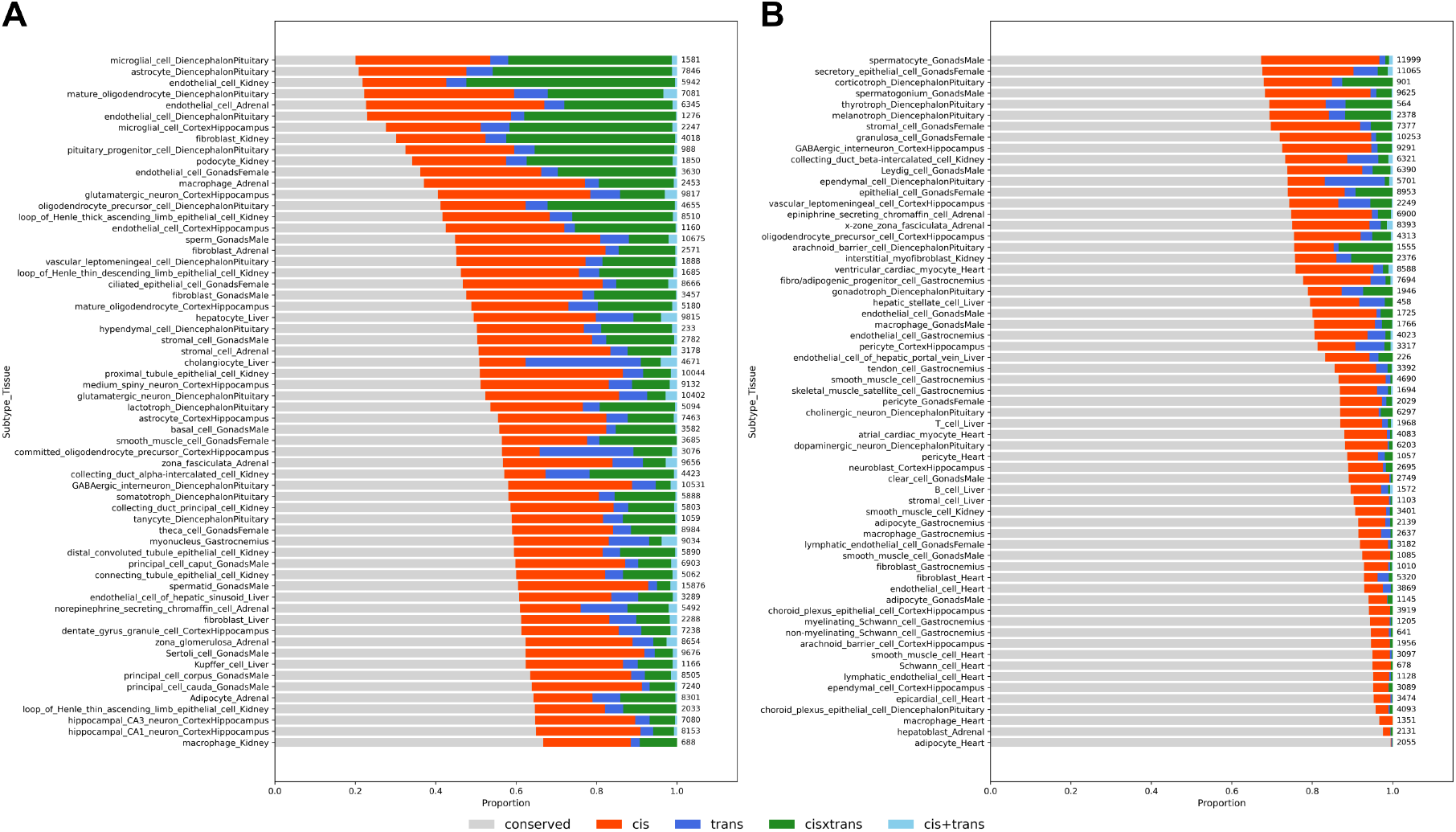
Regulatory composition in all cell-types in B6J-CAST trios. Proportion bar plots depicting regulatory composition across all cell types in B6J-CAST trios, ordered by proportion of conserved genes. (A) The 63 cell types with the highest and (B) the 63 with the lowest proportions of non-conserved genes.

**Figure S6.**
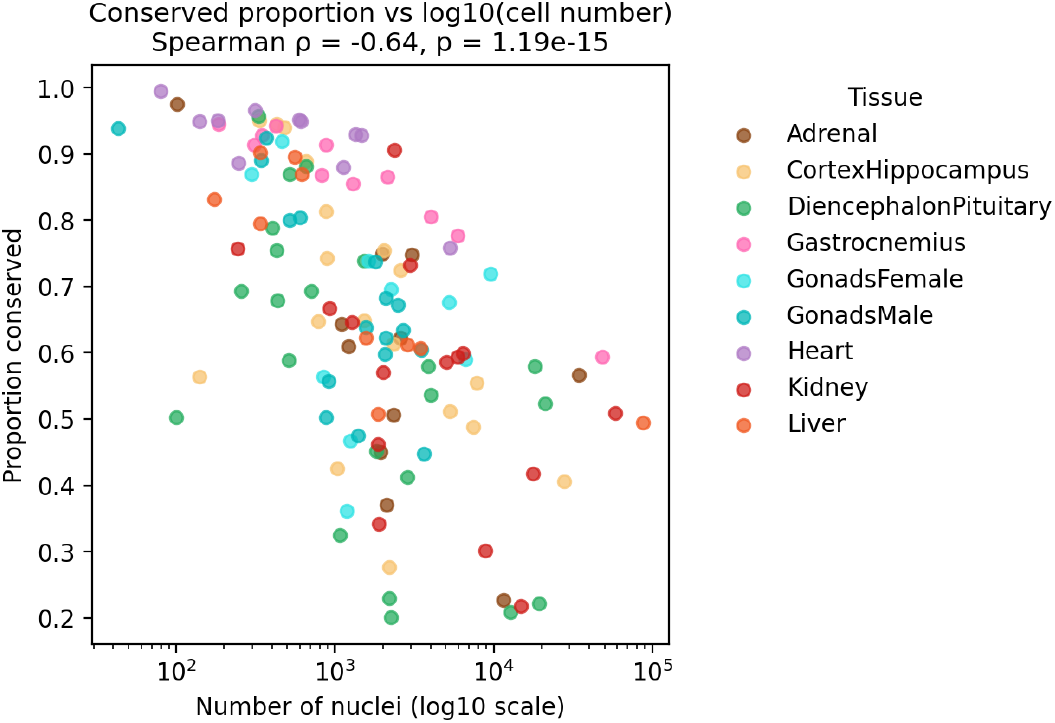
Relationship between number of nuclei in a cell type and proportion of genes identified as conserved in B6J-CAST trios. Scatterplot of proportion of genes conserved in each cell type by log10 number of nuclei in that cell type.

**Figure S7.**
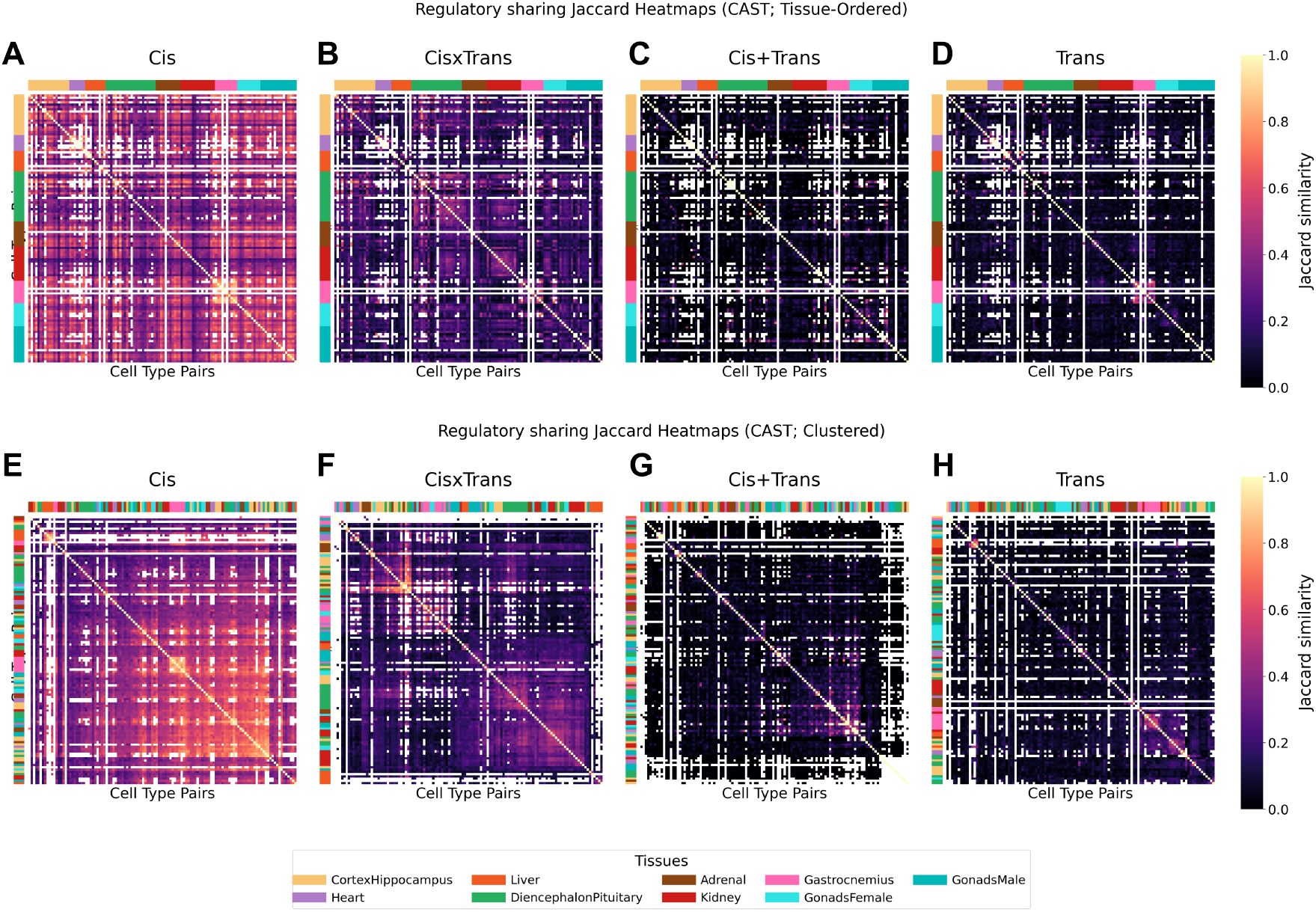
Enrichment of regulatory assignment sharing between pairs of cell types in B6J-CAST trios. (A-D) Heatmaps of regulatory sharing of cis-acting (A), cis*×*trans-acting (B), cis+trans-acting (C), and trans-acting regulatory effects (D), with cell types ordered by tissue. (E–H) Corresponding heatmaps with rows and columns hierarchically clustered. White cells (dropouts) indicate cell type pairs which share less than 50 non-conserved genes.

**Figure S8.**
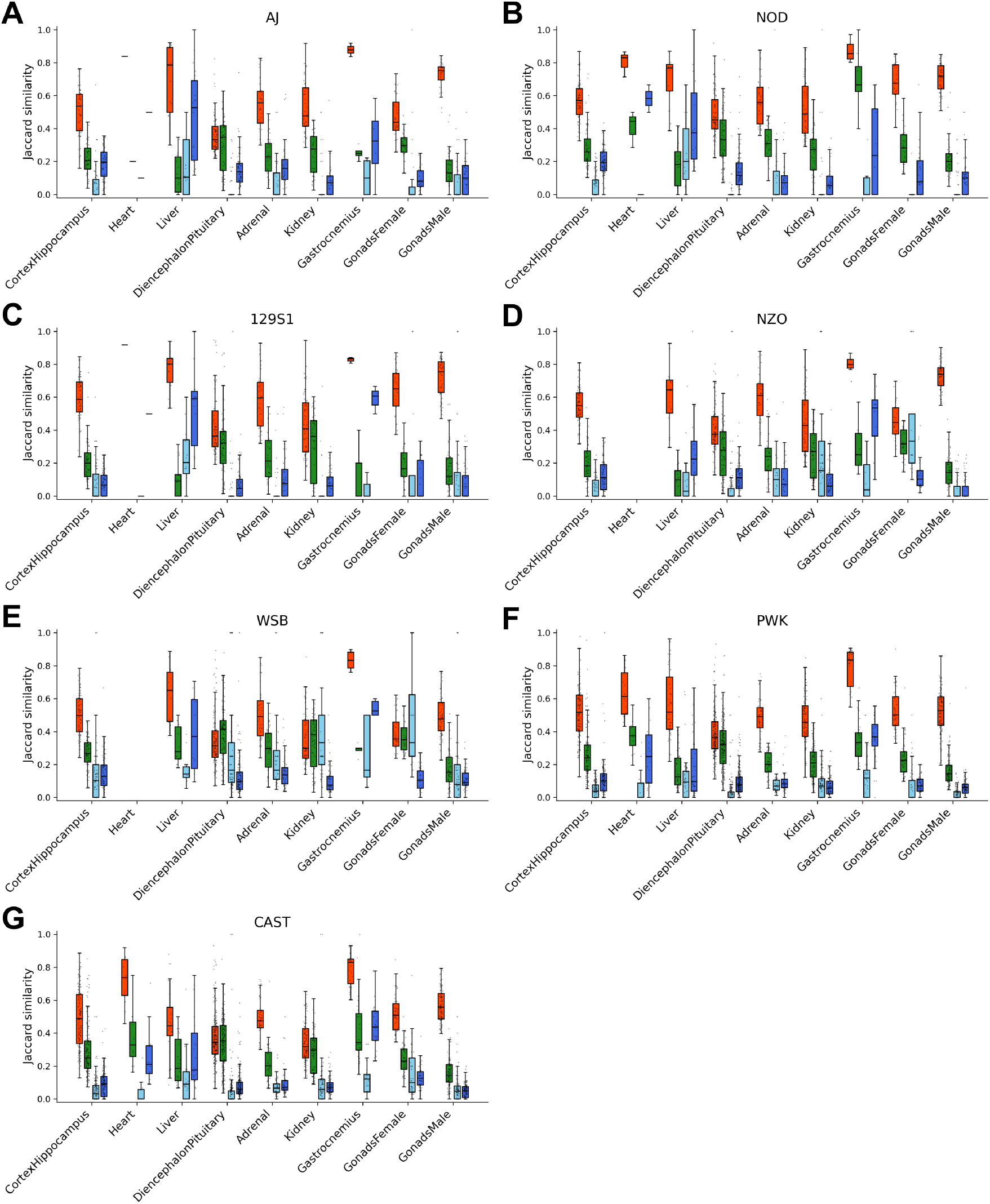
Tissue specific regulatory sharing across parental-F1 hybrid trios. (A) AJ, (B) NOD, (C) 129S1, (D) NZO, (E) WSB, (F) PWK, (G) CAST.

**Figure S9.**
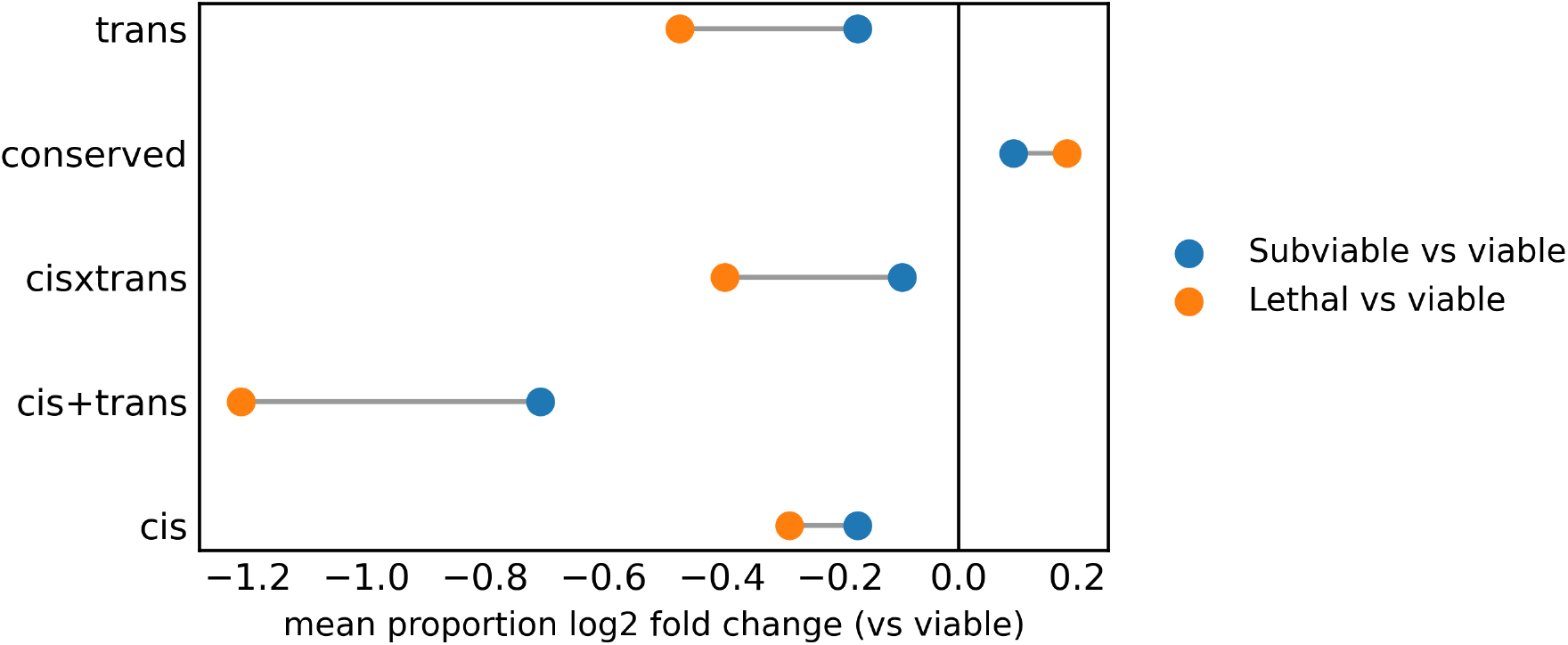
Shift in regulatory assignments associated with gene essentiality. Log2 fold change in median regulatory class proportions for subviable and lethal genes relative to viable genes across 126 cell-type–tissue combinations. Only genes observed in at least 33 combinations are included. See also Figure 3F.

**Figure S10.**
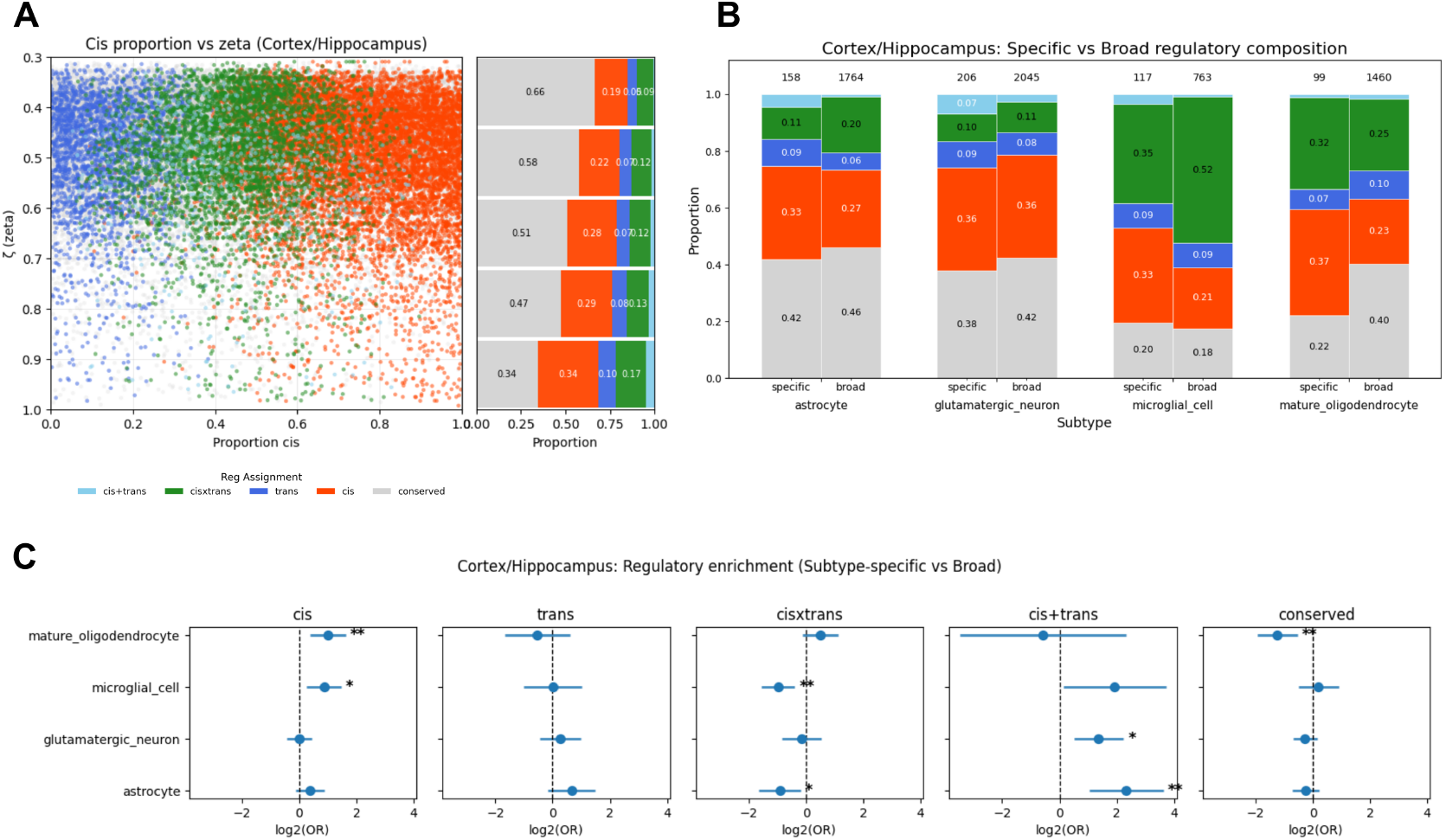
Relationship between cell type-specificity and regulatory assignments in cell types of the cortex and hippocampus in B6J-CAST trios. (A) Plot of proportion cis and *ζ* values (high *ζ* values indicate greater cell-type specificity) for genes expressed in cell types of the cortex and hippocampus. The proportion bar plot is divided into five bins based on *ζ* value. (B) Regulatory composition of cell-type-specific genes (*ψ*_block_ *>* 0.9) compared to broadly expressed genes across four representative cell types. (C) Enrichment of regulatory assignments in cell-type-specific genes relative to broadly expressed genes. Odds ratios (OR) and significance were calculated using a two-sided Fisher’s exact test to determine if specific regulatory classes (e.g., cis-acting) are overrepresented in cell-type-specific gene sets compared to the broadly expressed background (* q < 0.05, ** q < 0.01, *** q < 0.001).

